# The Brazilian population of *Fusarium oxysporum* f. sp. *cubense* is not structured by VCG or by geographic origin

**DOI:** 10.1101/2022.02.08.479520

**Authors:** Izabel C.A. Batista, Daniel W. Heck, Alessandro Santos, Gabriel Alves, Camila G. Ferro, Miguel Dita, Fernando Haddad, Sami J. Michereff, Kamilla C. Correia, Christiana F. B. da Silva, Eduardo S. G. Mizubuti

**Affiliations:** Departamento de Fitopatologia, Universidade Federal de Viçosa, MG 36570-900, Brazil; Alliance of Bioversity International and CIAT, Cali, Colombia; Embrapa mandioca e Fruticultura, BA, Brazil; Universidade Federal do Cariri, CE 63048-080, Brazil; Embrapa Agroindústria Tropical, CE, Brazil

**Keywords:** Panama disease, population biology, epidemiology

## Abstract

Fusarium wilt, caused by the soil-borne fungus *Fusarium oxysporum* f. sp. *cubense* (Foc), is considered one of the most destructive diseases of bananas. Paradoxically, knowledge of the genetics of the pathogen population in the Americas is very limited. In this study, a collection of 178 monosporic isolates from several banana producing regions, located in different climatic zones along a South to North transect in Brazil, was formed to assess the genetic structure of the population of Foc. The isolates underwent pathogenicity tests, PCR diagnosis for the detection of Tropical race 4 and screening to *SIX* homologs. The VCG of 119 isolates was determined by pairing against 17 testers. A group of 158 isolates was selected for microsatellite genotyping. There was moderate diversity of Foc in Brazil. Eight VCGs were identified: 0120, 0122, 0124, 0125, 0128, 01215, 01220, and 01222, of which 78% of isolates belong to a single VCG, while 22% of isolates belong to complexes of VCGs. The distribution of VCGs is uneven and independent of the banana genotype. VCGs were correlated with homologs of the *SIX* genes and varied according to geographic regions. Four SSR loci were polymorphic and on average 7.5 alleles were detected per locus. Thirty-five multilocus genotypes (MLGs) were identified. There was no association between VCG and MLGs and no genetic structure of the population of Foc in Brazil was detected.

## Introduction

Recent migratory events of the tropical race 4 (TR4) of *Fusarium oxysporum* f. sp. *cubense* (E. F. Smith) Snyder and Hansen (*Foc*), the ascomycete that causes Fusarium wilt of bananas (FWB), have been widely disclosed and have attracted the attention of the public at large. The pathogen variant, TR4, recently reported in Colombia and Peru, threatens the world’s largest banana exporter countries from Central and South America (García-Batidas et al. 2020; Acuña et al. 2021). There is an urgent need to establish mitigation plans to reduce the impacts of the introduction of TR4 in Central and South American countries. Such practices need a sound epidemiological basis that will certainly depend on both ecological and genetic information of the population of the pathogen.

Fusarium wilt of bananas is a destructive disease worldwide, regardless of the race of the pathogen (Dita et al. 2018). The fungus infects the banana plant through the roots, colonizes and moves in the vascular bundle, and blocks the transport of nutrients and water, causing a reddish-brown discoloration of the rhizome and pseudostem. As a consequence, leaves and whole plants normally collapse after FWB development (Stover 1962; Fan et al. 2017, Liu et al. 2020). After inspecting more than 90 ha during an extensive survey of commercial fields of banana in Brazil, the average incidence of FWB caused by non-TR4 races was approximately 11%, with estimated yield losses of 1.8 t.ha^−1^.year^−1^ (Heck et al. 2021).

The genetic variation and phenotypes of *Foc* are likely to affect the development of FWB epidemics in distinct ways. Specifically, there are variants of *Foc* in relation to the capacity to infect banana genotypes. To date, three races of *Foc* are recognized (Ploetz 2006): Race 1 (R1) is pathogenic to ‘Gros Michel’ (AAA), ‘Prata’ (‘Pome’) and ‘Maçã’ (‘Silk’) (AAB) varieties. Race 2 (R2) affects Bluggoe, a cooking banana group (ABB) and some tetraploids (AAAA) (Stover and Waite 1960; Ploetz 1990). Race 4 is subdivided into Tropical Race 4 (TR4), recently suggested to be classified as *F. odoratissimum* (Maryani et al. 2019; Torres-Bedoya et al. 2021), and Subtropical Race 4 (SR4). Both race 4 variants can cause FWB in cultivars belonging to the Cavendish subgroup (AAA), and all other banana varieties susceptible to R1 and R2. SR4 is known to affect Cavendish in areas subjected to low temperatures and other predisposing stress factors, while TR4 affects Cavendish in both tropical and subtropical conditions (Buddenhagen 2009).

In addition to the variability at the pathogenicity level that defines races, there is variation within the *formae speciales* regarding vegetative compatibility groups (VCGs). The recognition of VCGs within a morphologically homogeneous population allows the identification of potentially genetically isolated subpopulations with possible distinct characteristics, such as pathogenicity or geographical origin (Leslie 1993). At least 24 VCGs are reported in *Foc* (Fourie et al. 2009; O’Donnell et al. 2009; Mostert et al. 2017). Furthermore, some VCGs are compatible with each other and form complexes. Three VCG complexes are reported for *Foc*: VCG complex 0120/15, 0124/5/8/20/22 and 01213/16 (Mostert et al. 2017). Vegetative compatibility groups can play an important role in structuring the pathogen population, since nuclear exchange is more likely to occur among individuals of the same VCG. Nevertheless, there is no strict relationship between VCGs and races of *Foc*.

The use of molecular markers to assess genetic variability of the Foc revealed a population that is not structured according to host or region in Brazil. In a previous study, Costa et al. (2015) used the SSR markers to investigate the diversity of *Foc* in the states of Bahia, Ceará, Rio Grande do Norte, Minas Gerais, and Rio Grande do Sul. Although it was not possible to observe any population structure among isolates, this is the most detailed study on the genetic structure of *Foc* in the Americas to date. However, populations change over time and it would be interesting to quantify the amount of genetic variability present in the current Brazilian population using more makers and to try to identify any distribution of potential variants in the banana producing regions.

Additional efforts are now being directed towards the molecular detection of virulence-related genes to complement population analyses. In *F. oxysporum* f. sp. *lycopersici* (*Fol*) a group of cysteine-rich effectors that are secreted in xylem (*SIX*) are required for full virulence to its host and reside on an accessory chromosome, which can be transferred horizontally between strains (Zhao et al. 2020). The screening of the *SIX* genes in a population of *Foc* can provide information to make inferences about race and VCGs (Carvalhais et al. 2019). Thus, assessing the population composition regarding *SIX* genes can add another trait to estimate genetic variability of *Foc* in Brazil. To date, there is no information regarding the *SIX* gene-profile for populations of *Foc* in Central and South America.

The characterization of VCGs currently present in Brazil combined with the profiling of *SIX-*genes and fluorescence-based SSR markers to genotype individuals from different areas can add useful information about the population biology of *Foc*. This will contribute to a better understanding of the evolutionary processes that may be acting upon the pathogen populations and to help establish effective practices to the management of FWB. The objectives of this study were to investigate the presence of TR4 in a recent collection of isolates and to determine the genetic structure based on an expanded set of markers: *SIX* genes profiling, VCG test, IGS PCR-RFLP and SSR markers of *F*. *oxysporum* f. sp. *cubense* in a Brazilian population.

## Material and methods

### *Foc* isolates

Fungal isolates were sampled from different banana cultivars grown in distinct geographic regions in Brazil. Pseudostem tissue of FWB symptomatic plants or non-pure isolates were sent to the Departamento de Fitopatologia at the Universidade Federal de Viçosa (UFV). Monosporic cultures were obtained and mycelial discs from these cultures grown on synthetic low-nutrient agar (SNA; Nirenberg 1976) were maintained in microtubes at 10 °C.

Pathogenicity tests were performed for 212 isolates in a highly susceptible cultivar (‘Maçã’) to confirm the classification as *Fusarium oxysporum* f. sp. *cubense*. The methods used in the pathogenicity tests were described previously (Perez-Vicente et al. 2014). Population analyses were conducted with *Foc* isolates originated from different locations such as to include individuals from a wide range of edaphoclimatic conditions. Furthermore, the isolates used in the analyses were collected from fields planted with different cultivars and banana groups such as ‘Pome’, ‘Silk’, and ‘Cavendish’ located along a South to North transect in eastern Brazil (Table S1).

### DNA extraction

Mycelial discs of *Foc* colonies were transferred to Erlenmeyer containing potato dextrose (PD) medium (Leslie and Summerell, 2006) and grown for 7 days at 25 °C, in the dark. After this period, the mycelial mass on PD was transferred to sterilized filter paper and the mycelium retained on the filter was allowed to dry at room temperature for 24 h. The dried fungal mass was stored at −20 °C. DNA extractions were carried out using the Wizard® Genomic DNA Purification Kit (Promega, Madison, WI, United States of America) following Lehner et al. 2017. The extracted DNA was quantified by spectrophotometry (Nanodrop 2000), and its integrity assessed by electrophoresis on 1% agarose gel. The DNA was diluted to the final working concentration of 10 ng /μL^−1^.

### *Foc* and TR4 PCR diagnostic

The 212 *Fusarium oxysporum* isolates were screened to confirm the identification of *formae speciales* “cubense” and for the potential occurrence of the variant TR4. The screening was conducted using the R4 specific PCR (Lin et al. 2009), TR4 specific PCR (Dita et al. 2010), and *Foc*-specific PCR developed to amplify a conserved region of 122 bp of the genome of *Foc* (Heck et al. *unpublished data*). PCR reactions were combined in a multiplex reaction herein developed (M1; Table S2). The multiplex reactions were performed in a volume of 12.5 μL, which consisted of 2.5 μL of 5x GoTaq® Reaction Buffer (Promega), 0.25 μL of dNTP (10 mM), 0.2 μL of the forward and reverse primers each (1 μΜ), 0.05 μL GoTaq® DNA Polymerase (5u/μL), 1 μL template DNA (~ 10 ng/μL), and Milli-Q water to 12.5 μL. The multiplex-PCR conditions were: initial denaturation for 4 min at 95 °C, followed by 35 cycles of 30 sec denaturation at 95 °C, annealing for 1 min at 58 °C and extension for 1 min at 72 °C, with a final extension of 10 min at 72 °C. The presence or absence of bands was solved in an electrophoresis analysis using a 1.5% agarose gel. The following analyses were performed in pathogenic isolates and positive for the *Foc*-specific PCR analyses.

### *SIX* genes profiling

A subset of the isolates classified as *Foc* (N = 143) were subjected to PCR test based on the amplification of *SIX* genes homologs using the primer sets and multiplex and individual PCRs. Sixteen *SIX* genes (*SIX*1 through *SIX*14) were analyzed, including the homologues *Foc*-*SIX*8a and *Foc*-*SIX*8b, which are candidate molecular markers that allow differentiation of SR4 and TR4, in addition of race 4 from races 1 and 2 (Table S2). The 16 primer-pairs were distributed in four multiplex and two singleplex reactions. The multiplex PCR was performed in a 12.5 μl volume containing 2.5 μL of 5x GoTaq® Reaction Buffer (Promega), 0.25 μL of dNTP (10 mM), variable concentrations of primers (Table S2), 0.05 μL GoTaq® DNA Polymerase (5u/μL), 1 μL template DNA (~ 10 ng/μL), and Milli-Q water to 12.5 μL. The reactions were conducted in a T100 thermal cycler (BioRad Laboratories Inc., Hercules, CA, United States of America) using the conditions described in the previous section. The multiplex PCR results were analyzed using electrophoresis on a 1.5% agarose gel and expressed as the presence/absence for each of 14 *SIX* genes in addition to the homologues *Foc*-SIX8a and *Foc*-SIX8b. Clustering analysis based on presence/absence data was performed using the UPGMA algorithm in the *stats* R package (R Core Team, 2020).

### Vegetative compatibility groups

Vegetative compatibility tests were conducted to determine the VCG group of *Foc* isolates. Wild type strains were first grown on minimal medium (MM) to check for dense mycelial growth. After 3-5 days, when the colonies had grown vigorously, mycelial discs from each isolate were placed on plates containing chlorate medium (MM + 1.5 to 4.0% KClO_3_) and kept at 25 °C for 7-21 days (Puhalla 1985).

Mycelial discs from spontaneous KClO_3_-resistant sectors that did not form aerial mycelium were classified as *nit* mutant and were phenotyped as *nit*1, *nit*3 or *nit*M mutants based on their inability to assimilate nitrogenous compounds on four phenotyping media MM supplemented with NaNO_3_ (2 g/L), NaNO_2_ (0.5 g/L, Sigma Chemical Co., St. Louis), hypoxanthine (0.2 g/L, Sigma Chemical Co.), and ammonium tartrate (1.6 g/L). The pH of the culture media was adjusted to 6.5 using NaOH 1 M (Correll et al. 1987).

Strains of 17 VCGs testers were sent by Prof. Randy Ploetz of the University of Florida, USA (Table S3). Due to quarantine restrictions, tester 01213/16, associated with TR4, was not available for the assays. The testers (VCGs 0120, 0121, 0122, 0123, 0124, 0125, 0126, 0128, 0129, 01210, 01211, 01212, 01214, 01215, 01219, 01220, and 01222) were paired with the assigned *Foc* isolates on MM supplemented with NaNO_3_ as the sole nitrogen source. The *nit*1 mutants were paired with its *nit*M mutant cognate to confirm heterokaryon self-compatibility, and then with the *nit*M or *nit*1 of the tester to determine VCG identity (Leslie 1993; Leslie and Summerell 2006). Pairing tests on Petri dishes were kept at 25 °C in complete darkness and scored for heterokaryon formation after 14 days. Complementation tests were repeated at least twice. Positive reactions were considered those that presented dense aerial mycelium growth in the line of intersection between paired mutants, confirming the classification of these strains as belonging to the same VCG. Negative reactions were those that had incipient growth in the line of intersection between the paired mutants.

### Clade assignment based on IGS PCR-RFLP

A fragment of the IGS region of rDNA was amplified using primers PNFo (5’-CCCGCCTGGCTGCGTCCGACTC - 3’) and PN22 (5’- CAAGCATATGACTACTGGC - 3’) (Edel et al. 1995). PCRs were performed with 10 ng of template DNA in 50 μl reaction volume using 0.5 μl of each primer; 0.5 μl of dNTP; 0.3 μl of Taq DNA polymerase; and T5x Buffer (Promega, Madison, USA). PCR conditions were initial denaturation of 2 min of 95 °C, 30 cycles of 90 s at 95 °C, 60 s at 50 °C, and 90 s at 72 °C, with a final extension of 5 min at 72 °C. The amplification products were visualized by electrophoresis on a 0.8 % agarose gel stained with Gel Red (Biotium, Hayward, USA) at 80 V.

Five restriction enzymes were used for PCR-RFLP analysis: *Ava*I, *Bbv*I, *BceA*I, *BsrD*I, and *CviQ*I (New England BioLabs). All enzymes were used separately in PCR-RFLP digestion reactions, which was performed in a total volume of 20 μl and consisted of 5 μl IGS PCR product, 2U of restriction enzyme and 2 μl of the provided restriction buffer. After incubation at 37 °C for 1 h or 3 h, depending on the manufacturer recommendation, the restricted fragments were visualized on a 3% agarose gel electrophoresis at 60 V for 180 minutes (Fourie et al. 2009).

The pattern of digestion with *Ava*I enzyme allowed to differentiate *Foc* isolates into clades A and B (Fourie et al. 2009). Clade A isolates were then digested with the following enzymes: *BceA*I to separate the *Foc* lineage V from lineages I, II, III and IV; *CviQ*I to separate lineages I and II from lineages III and IV; and *BsrD*I to separate lineage III and lineage IV. Isolates of clade B were digested with the enzyme *Bbv*I to separate lineage VII from lineages VI and VIII (Fourie et al. 2009).

### Microsatellites genotyping

The genetic variability of the population was analyzed with nine microsatellite markers described by Bogale et al. (2005). Forward primers were labeled with fluorophores 6-FAM, VIC, NED or PET (Table S4). The primers were first tested in an individual PCR for optimization. Only primers producing good quality PCR products for all samples were retained for further analysis. The reactions were performed using the Multiplex 5X Master Mix (New England Biolabs, Inc.) in a final volume of 12.5 μl following the protocol described by the manufacturer. The final concentration of the primers was 0.2 μM. The PCR products were lyophilized and fragment analysis was done using the GeneScan-500 LIZ size standard (Applied Biosystems) on an ABI 3730xl analyzer. The fragments were analyzed with the software GENEMARKER v.1.191 (SoftGenetics). The size of DNA fragments was manually binned into alleles according to the number of repeat units at each locus.

### Data analyses

The *poppr* R package (Kamvar et al. 2014) was used to calculate gene diversity (Nei 1978) by locus, the number of multilocus genotypes (MLGs), and the index of association (r_d_). Diversity indices as genotype richness, the exponential of Shannon’s entropy, and the inverse of the Simpson’s concentration were estimated by the Hill numbers (N) of orders 0, 1, and 2 for the whole population. The diversity analyses were conducted with the iNEXT package (Hsieh et al. 2016). A principal components analysis (PCA) was performed on the data set without any population assignment to identify clusters of individuals. The *adegenet* R package (Jombart 2008; Jombart and Ahmed 2011) was used to create a minimum spanning network based on Bruvo’s genetic distance (Kamvar et al. 2014; 2015).

## Results

A total of 178 monosporic isolates were selected for this study. Selection criteria were: different locations from a North-South transect in Brazil, and banana genotypes. Only *Fusarium oxysporum* f. sp. *cubense* isolates were used. These isolates were chosen based on the pathogenicity test and on the PCR diagnostic using a set of primers that is able to detect *Foc* strains (Heck et al. *unpublished data*). A subset of 140 isolates were selected for the VCG test and *SIX* gene profiling. The microsatellite genotyping was performed for 158 isolates. A total of 98 isolates were subjected to both the SSR genotyping and VCG test (Table S1).

### TR4 PCR-based diagnostic

None of the Brazilian isolates tested were positive for the TR4 variant. Despite recent claims that this method can generate false positives during the diagnostic process for TR4, there are no reports of false negatives (Magdama et al. 2019). Therefore, this result suggests that Brazil may still be a free area of TR4. The same isolates were subjected to a screening using the R4 primers which amplifies for both SR4 and TR4 strains. From those, 74 of 178 (41.6%) isolates tested were positive for race 4. However, as TR4 was not detected in any of these isolates, the positive isolates were classified as SR4 strains.

### *SIX* genes profiling

*SIX1*, *SIX7*, *SIX8* and *SIX13* genes were most frequent (>42.8%) among the *Foc* isolates examined in this study (Table S1). *SIX1* were detected in all 140 isolates, followed by *SIX8* detected in 86 isolates, *SIX13* detected in 73 isolates and *SIX7* detected in 60 isolates. Additionally, the *SIX12* gene was detected in three isolates and the *SIX2* gene was detected in only one isolate. The homologues of *SIX8* genes identified (*Foc-SIX8a and Foc-SIX8b*), proposed to differentiate race 4 from race 1 and race 2, also revealed through a unique combination that TR4 is not present. Since in previous studies *Foc*-SIX8b was present in all SR4 isolates and was not detected in TR4 isolates.

For the SIX genes detected among the *Foc* isolates, two major clusters were formed in the dendrogram based on vegetative compatibility (Figure 2). Cluster 1 contained isolates belonging to VCGs 0120, 01215, its complex 0120/01215, and a low frequency (7.2%) of unknown VCGs. In cluster 1, 92.8% of the isolates originated from the South and Southeast regions. Cluster 2 contained isolates from different arrangements of the VCGs tested, including 0120, 0122, and the complexes formed by 0124/5/8/20/22. In this cluster, neither VCG 01215 nor its complex 0120/15 were observed. Isolates of unknown VCG were predominantly (89.4% of total unknown VCG isolates) grouped in cluster 2. Finally, 81.8% of these isolates were collected from banana plantations located in the North and Northeast regions of Brazil.

### Vegetatitive compatibility group

Out of 140 isolates, *nit* mutants were generated for 119 isolates (85%). Eight isolates were chlorate resistant (*crn* mutants) and could not be used in VCG testing. Most isolates (N = 88) had both *nit*1 and *nit*M or *nit*1 and *nit*3 mutants, and no self-incompatible mutant was noticed. When the 119 *Foc* mutants were paired with the 17 testers of known VCGs available, 46 isolates did not form heterokaryon with any of the testers, 73 isolates formed heterokaryons, of which 57 isolates belong to a single VCGs (78%) and 16 isolates belong to complexes of VCGs (22%). Eight of the 24 known VCGs were identified in Brazil: 0120, 0122, 0124, 0125, 0128, 01215, 01220 and 01222.

Among the Brazilian *Foc* isolates that belong to a single VCG, the most frequent was VCG 0120 (69.9% of the isolates), followed by the VCG 0124 (4.1%), VCG 0122 (1.4%), VCG 01215 (1.4%) and VCG 01222 (1.4%). In the group of isolates that belong to complexes of VCGs, 12.3% of isolates belong to complex VCG 0120/15, while 9.6% of isolates belong to VCG 0124/5/8/20/22. For complexes involving many VCGs, e.g. 0124/5/8/20/22, the complementation of the isolates was not always detected for all its components. In this study four VCG combinations were detected: 0124/5/8/22, 0124/5/22, 0125/8/20, and 0124/22. No isolate 0124/5/8/20/22 was found in our analyses. For this reason, these new arrangements (0124/5/8/22, 0124/5/22, 0125/8/20, and 0124/22) were counted as independent VCGs and not as belonging to a single VCG 0124/5/8/20/22.

Highest VCG richness was recorded in Ceará state (N = 8 isolates), located in the Northeast region (Figure 1B). Six VCGs were detected in Ceará: 0122, 0124, 0125, 0128, 01220 and 01222. Some of these VCGs formed specific arrangements of VCG complexes: 0124/22, 0124/5/22, 0124/5/8/22 and 0125/8/20. All *Foc* strains from Ceará were isolated from the Pome subgroup (AAB). Four VCGs were found in São Paulo state (N = 36): 0120, 0120/15 and 0124/22; three in Minas Gerais (N = 36): 0120, 01215 and 0124; and three in Maranhão state (N = 3), all as VCG complex 0124/5/22. The isolates from São Paulo and Minas Gerais were sampled mainly from the Pome subgroup (AAB). In the states of Santa Catarina (N=20), Paraná (N=7), Bahia (N=29) and Rio Grande do Sul (N=4), only VCG 0120 and/or its complex 0120/15 were found. In the state of Pernambuco (N=3) only VCG 0124 was found. Therefore, no relationship between VCGs and banana cultivar or genotypes was observed.

**Figure 1.**
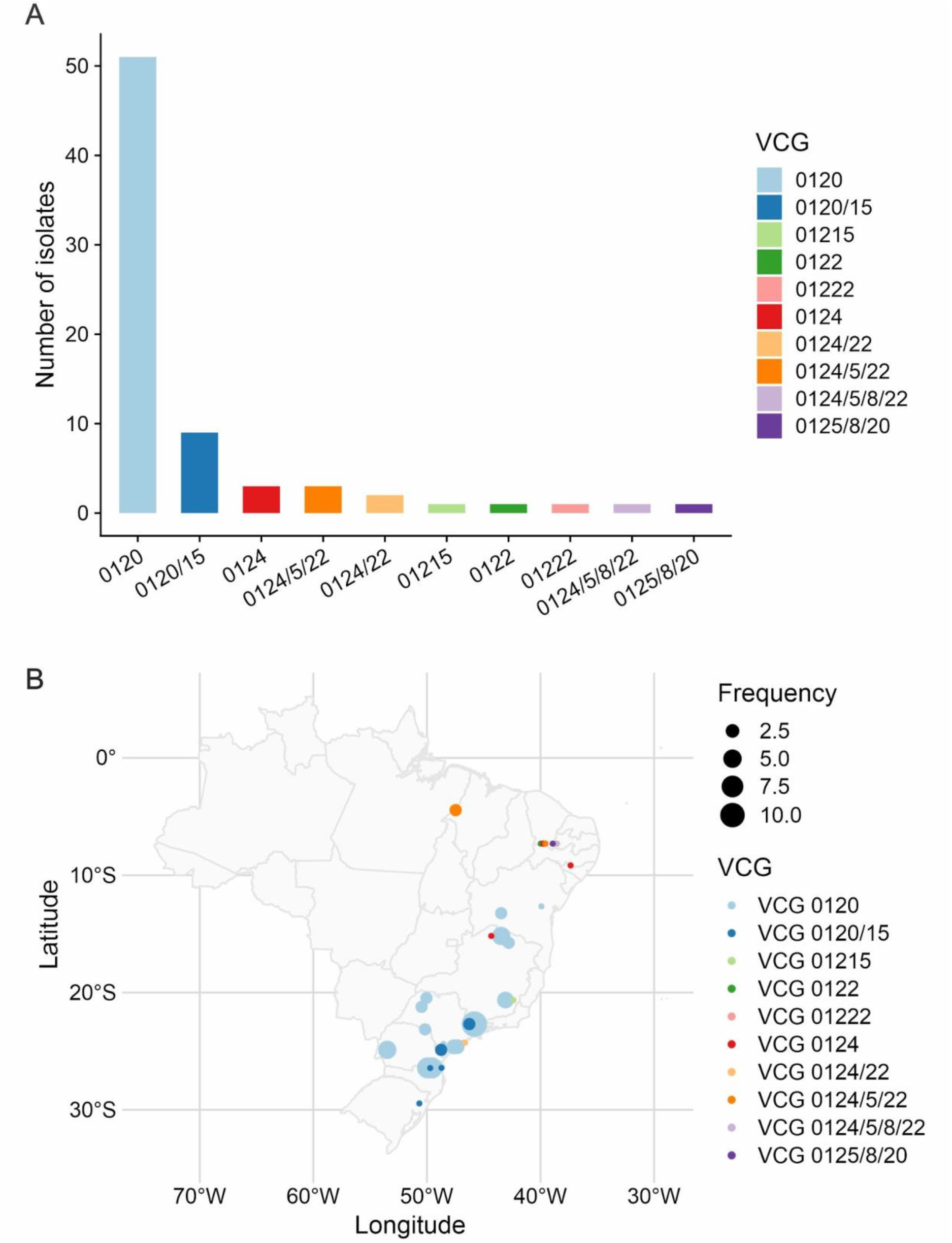
A. Vegetative compatibility groups (VCGs) of *Fusarium oxysporum* f. sp. *cubense* (Foc) occurring in Brazil. B. Distribution of the Foc VCGs found in Brazil. Jitter function was used while plotting the points. Thus, there are in slight distortions in the representation on the map.

**Figure 2.**
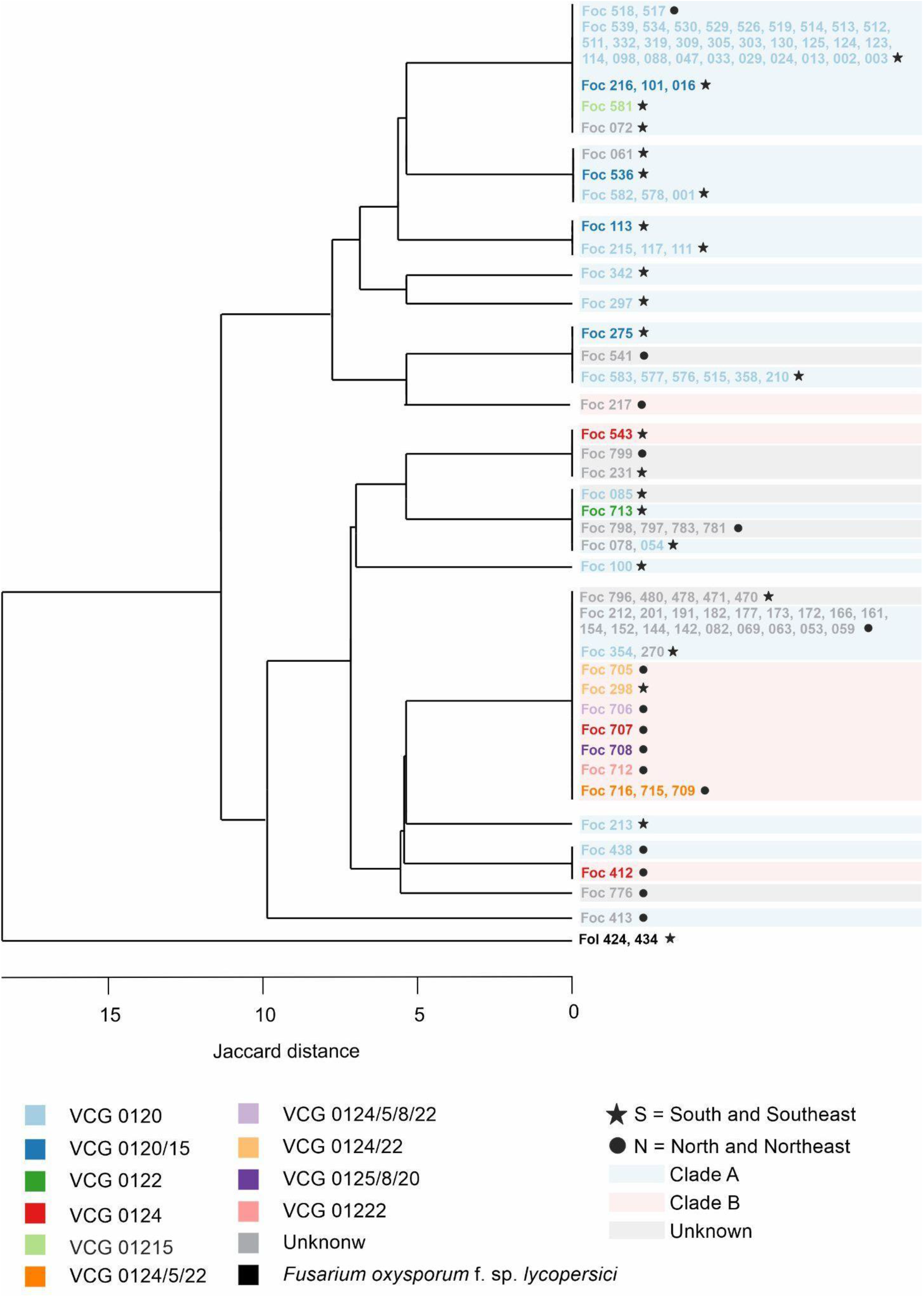
Dendrogram generated using the unweighted pair-group method with arithmetic means (UPGMA) of Jaccard distance coefficients based on presence/absence of *SIX* genes in 157 isolates of *Fusarium oxysporum* f.sp. *cubense* sampled from different regions and banana cultivars. Different colors refer to the vegetative compatibility group of isolates. Terminal areas shaded with different colors correspond to clades A, B, or unknown.

To answer if there was a significant relationship between the occurrence of some VCGs and the geographic location, a Fisher’s exact test was conducted. The frequency of the two prevalent VCGs in Brazil, 0120 and 0124, varied (*P* < 0.001) among regions. There was uneven distribution among North-Northeast and South-Southeast regions. The VCG 0120 and its complex with 01215 (0120/15) are almost exclusively (96.6%) observed in the South and Southeast regions. The VCG 0124 and its complexes with 0124/5/8/20/22 were predominantly (81.8%) observed in the North and Northeast regions of Brazil.

### Clade assignment based on IGS PCR-RFLP

The IGS primers PNFo and PN22 amplified a 1700 bp fragment for all 140 *Foc* isolates. Based on the digestion with *AvaI* 96 isolates were classified as belonging to Clade A (68.5%) and 19 isolates to Clade B (13.5%). The PCR-RFLP profile for 25 isolates (18%) did not allow them to fit into either clade. The cleavage with the enzymes *BceA*I and *Bbv*I generated indiscernible patterns, preventing the classification of *Foc* isolates into lineages.

### Microsatellite genotyping

There was no amplification for four SSR loci: FO5, FO9, FO10, and FO13. An *in silico* analysis was performed to diagnose this issue and there were no hits or similarities between the regions of the microsatellite FO5, FO9 and FO10 (Genbank: AY931030, AY931029, and AY931028, respectively) against the database of four genomes of *Foc* assembled at the scaffold level (Genbank: AGND01, AMGP01, AMGQ01 and MBFV01). For the FO13 locus, although there were hits between the flanking sequence and the *Foc* genome, the software Primer3 (version 4.1.0) did not detect a match between the reverse primer (5’-3’: CTAAGCCTGCTACACCCTCG) and the sequence at a supposed locus. Thus, these four loci were not used for genotyping. All analyses were conducted based on the polymorphism found in five loci: FO2, FO11, FO14, FO17, and FO18.

A total of 35 genotypes were identified among the 158 isolates. The number of alleles at each locus varied from 2 (FO14) to 10 (FO2 and FO17). Two alleles were revealed at FO14 locus, but one occurred at low frequency (0.01), thus it was considered as uninformative and removed from the analyses. The four remaining loci had an average of 7.5 alleles per locus. The average gene diversity over all loci was 0.59, but estimates varied from 0.42 (FO11) to 0.67 (FO2). The sample size for the whole population was 158 individuals, and the genotypic diversity estimated by Hill numbers of orders 0, 1, and 2 were 35, 10.3, and 3.9 respectively (Table 1). Both, I_A_ and r_d_, calculated for clone-corrected dataset were significantly different from zero (*P* < 0.01) in the overall population. Thus, the random mating hypothesis was rejected, indicating a clonal population.

**Table 1:**
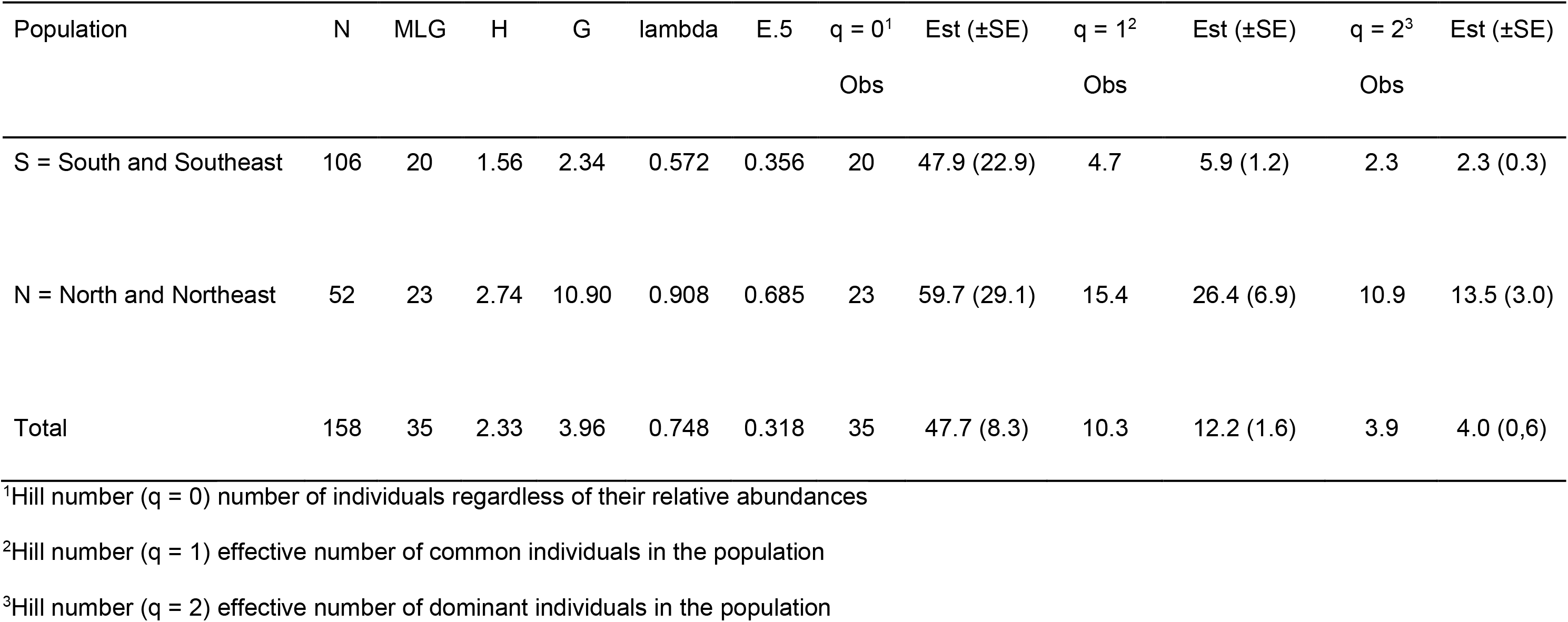
Summary of poppr index and observed (Obs) and estimated diversities (Est) and standard errors (SE) for the SSR genotyping data.

| Population | N | MLG | H | G | lambda | E.5 | q = 0<sup>1</sup> | Est (±SE) | q = 1<sup>2</sup> | Est (±SE) | q = 2<sup>3</sup> | Est (±SE) |
| --- | --- | --- | --- | --- | --- | --- | --- | --- | --- | --- | --- | --- |
|  |  |  |  |  |  |  | Obs |  | Obs |  | Obs |  |
| S = South and Southeast | 106 | 20 | 1.56 | 2.34 | 0.572 | 0.356 | 20 | 47.9 (22.9) | 4.7 | 5.9 (1.2) | 2.3 | 2.3 (0.3) |
| N = North and Northeast | 52 | 23 | 2.74 | 10.90 | 0.908 | 0.685 | 23 | 59.7 (29.1) | 15.4 | 26.4 (6.9) | 10.9 | 13.5 (3.0) |
| Total | 158 | 35 | 2.33 | 3.96 | 0.748 | 0.318 | 35 | 47.7 (8.3) | 10.3 | 12.2 (1.6) | 3.9 | 4.0 (0.6) |

<sup>1</sup>Hill number (q = 0) number of individuals regardless of their relative abundances
<sup>2</sup>Hill number (q = 1) effective number of common individuals in the population
<sup>3</sup>Hill number (q = 2) effective number of dominant individuals in the population

The PCA for all 158 isolates of the dataset indicated by the 35 MLGs was assessed grouping isolates according to location: N = North and Northeast (n = 52 / MLG = 23); S = South and Southeast (n = 106 / MLG = 20) (Table 1). The first two components were not sufficient to explain the total variation observed. The first component explained 18% of the total variation, and the second component 11.1% (Figure 3). No clear clusters of isolates could be identified. The formation of putative groups within quadrants was exploited through discriminant analysis of principal components (*data not shown*) and no biological or environmental pattern was noticed in the analyses. Thus, there was no evidence of population structure.

**Figure 3.**
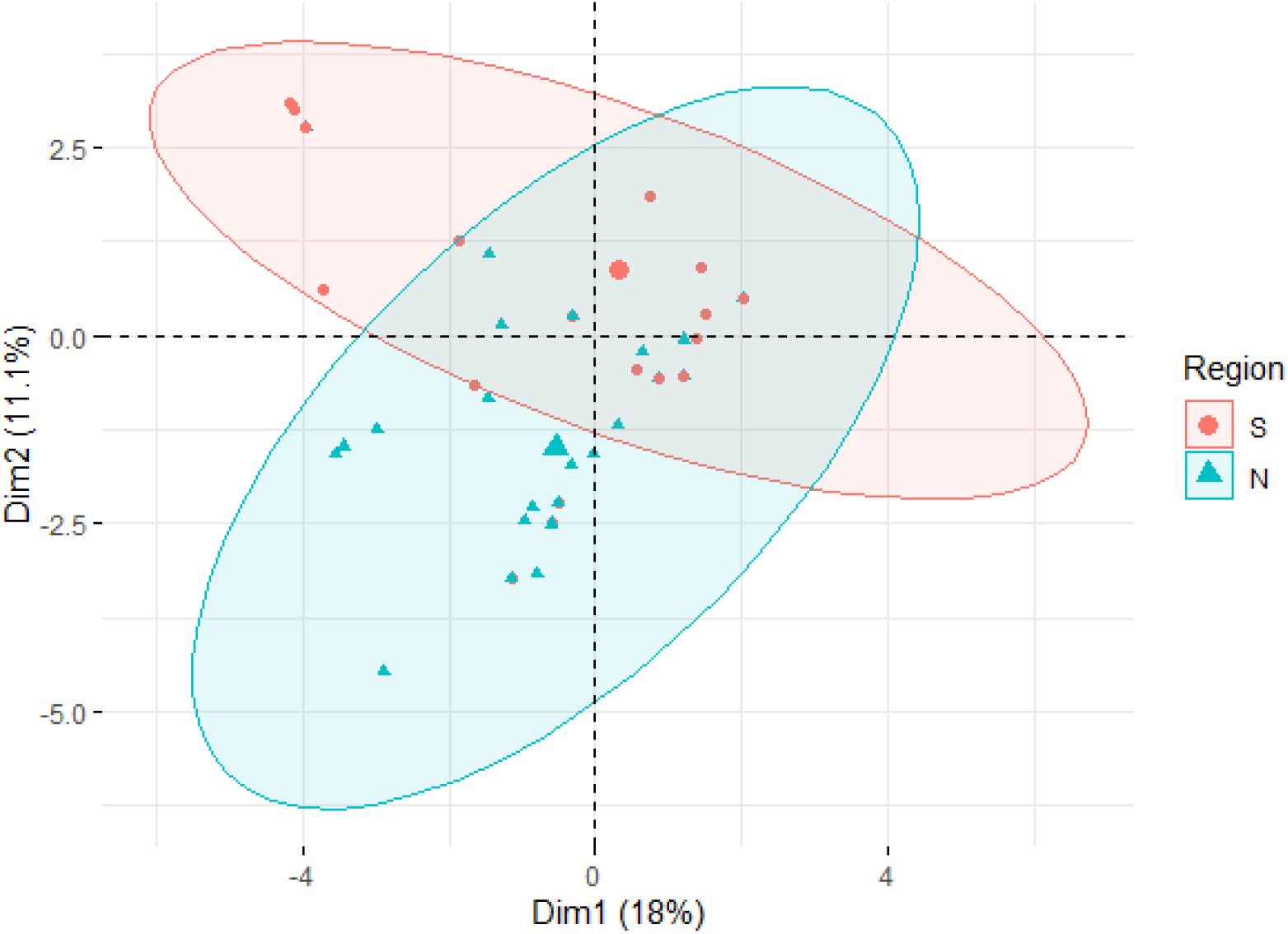
Scatter plot of principal component analysis (PCA) of 35 genotypes of *Fusarium oxysporum* f.sp. *cubense* sampled in different geographic regions (symbols) in Brazil. S: South and Southeast regions; N: North and Northeast regions.

The minimum spanning network also revealed no distinct pattern of genotype by VCGs or by location, with different VCGs within the same genotypes (Figure 4A), and genotypes from the North and Northeast contributing to the main groups of MLGs found in *Foc* (Figure 4B).

**Figure 4.**
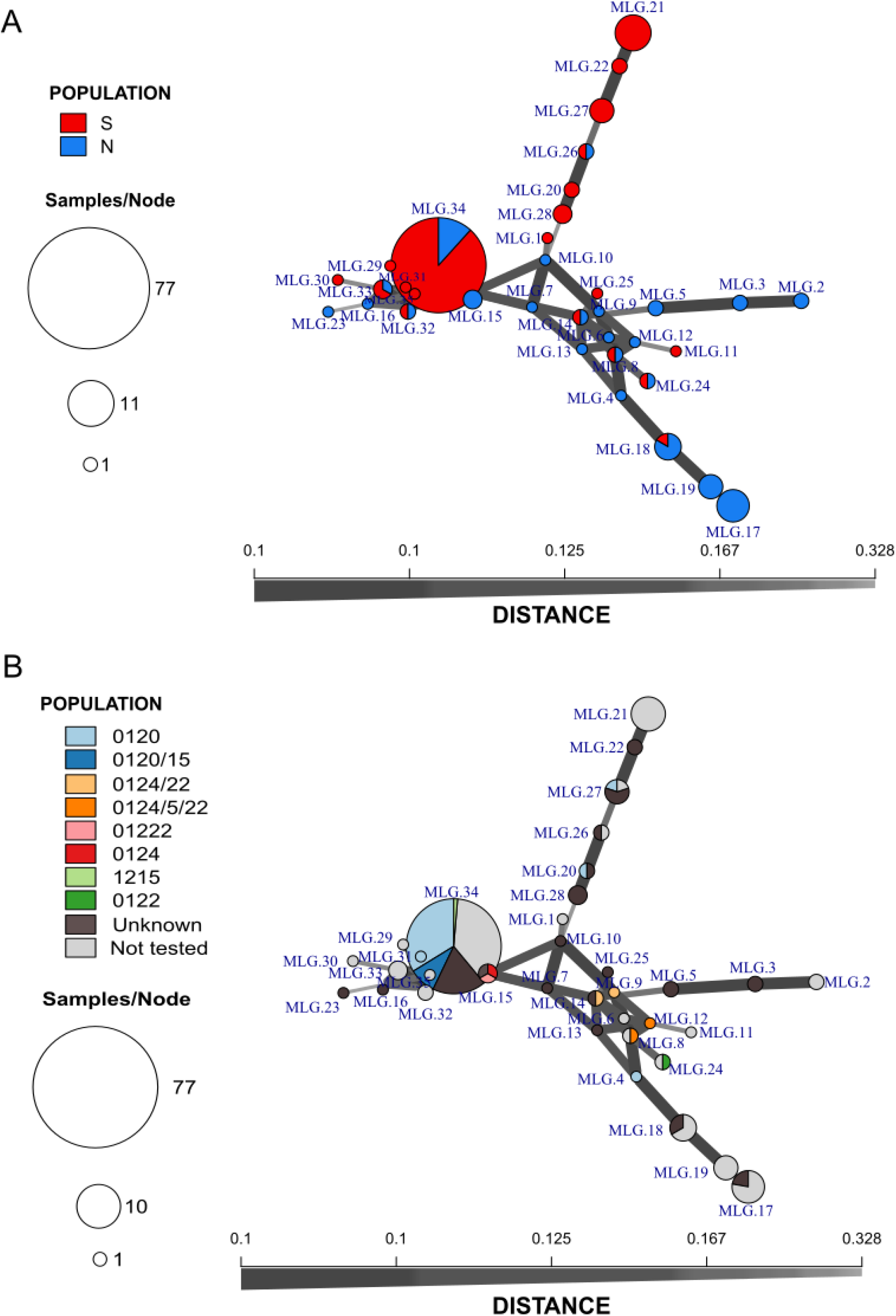
Minimum spanning tree of 35 multilocus genotypes detected in 158 isolates of *Fusarium oxysporum* f.sp. *cubense* according to A. geographic origin of the isolates; and B. the vegetative compatibility group.

## Discussion

In this study phenotypic and genotypic markers were combined to investigate the genetic structure of the population of *Foc* in Brazil. It was hypothesised that the population of *Foc* was highly variable regarding compatibility groups, since eight VCGs were found in a previous study that analyzed 44 isolates (Matos et al. 2009). In the present study, using approximately three times as many isolates, the same number of VCGs (8) was identified, but there were differences regarding them. The VCGs 0123, 0129 and 01210 were detected by Matos et al. (2009), but were not observed in the present study. In contrast, we found VCGs 0122, 01220 and 01222. To our knowledge, this is the first report of the occurrence of these VCGs in the Americas. Differences in the occurrence of VCGs could be due to true variation in the population or to variation in the set of testers used in the assays. We used a higher number of testers than in the work of Matos et al. (2009), but the same number of VCGs was detected. However, at least two major complexes of VCGs are reported in Brazil for the first time and no complex was reported in the previous investigation (Matos et al., 2009). Most likely, changes in the population of *Foc* resulted in different VCGs being detected in the present study.

Overall, VCG diversity in Brazil can be interpreted as high. Mostert et al. (2017), found 17 *Foc* VCGs from a total of 615 *Fusarium* isolates distributed in Asian countries, the putative center of origin of both banana and the pathogen. Eleven of these VCGs occur in Indonesia and Malaysia, when 47 and 67 *Foc* isolates, respectively, were phenotyped. In Brazil, there was greater phenotypic richness than in some Asian countries: Thailand (7 VCGs from 117 isolates), Bangladesh (6 VCGs from 61 isolates), India, Vietnam and Taiwan (5 VCGs from 66, 35 and 102 isolates) and Sri Lanka (4 VCGs from 27 isolates) (Mostert et al. 2017).

Highest VCG richness was in Ceará state. The high diversity of VCGs and the occurrence of different multilocus genotypes in Ceará state, may be related to the local evolution of the pathogen. Similar observations have been made in previous studies, in which the diversity of *Fusarium oxysporum* would be explained by the relation with local non-pathogenic isolates of *F. oxysporum* from nearby uncultivated areas and other *formae speciales* (Skovgaard et al. 2002; Fourie et al. 2009). Local pathogenic variants can be affected by different evolutionary mechanisms and as a consequence, a variable population originates in a region (Summerell et al. 2010). Apparently, local variants present in Ceará state can infect a specific banana cultivar, such as ‘Prata’ in Northeast Brazil, and contribute to high levels of haplotypic diversity in the region. The Ceará isolates originated from samples collected from small subsistence farms and rural backyard areas. Both sources were characterized by non-commercial, low-input areas with high diversity of crops and of propagative materials of unknown origin.

The occurrence of complex VCGs may allow plasmogamy among different lineages and potentially to a more variable population of *Foc*. Karangwa et al. (2018) have described a VCG complex as a set of individual VCGs that, in some cases, match each other. In this study all eight “single” VCGs, but VCG 0122, formed complexes. The VCGs 0120 and 01215 were also found as 0120/15 complex and the different arrangements of the VCG 0124/5/8/20/22.

The distribution of VCGs in Brazil depends on the region, and seems more related to the climatic conditions of the producing area than to the banana variety. The most prevalent VCG 0120 does not occur frequently in the North and Northeast regions, the hottest regions in Brazil where the average annual temperature varies between 25 °C to 36 °C (INMET, 2019). VCG 0120 was mainly found in the Southeast region, where the average annual temperature varies from 22 °C to 24 °C (INMET, 2019); Other VCGs and complexes that belong to Clade B were limited to a very specific region, such as 0122, 0125/8/20 and 0124/5/22 found only in the Northeast of Brazil. In all cases aforementioned, the isolates were sampled mainly from cultivar Prata (AAB - ‘Pome’). Contrary to what seems to happen in Brazil, in Asian countries the occurrence and distribution of VCGs seem to be influenced by banana varieties (Mostert et al. 2017). The higher diversity of banana genotypes in Asian countries could allow population structuring by host preference of *Foc*.

Another important note about the VCG 0120, is that many authors refer to it as the SR4 - subtropical race 4 (Buddenhagen, 2009; Czilowski et al. 2017). The lack of accurate markers for the detection of *Foc* races precludes this association. However, some strains of VCG 0120 characterized in this study show the STR4 phenotype, since they were collected from symptomatic Nanica and Nanicão cultivars (Cavendish subgroup - AAA) sampled from São Paulo and Santa Catarina states, located in the Southeast and South regions of Brazil, respectively. Both states can experience low temperatures during the winter, varying from 16 to 22 °C in Santa Catarina, and from 18 to 24 °C in São Paulo (INMET, 2019). The occurrence or predominance of VCGs in different geographic locations in Brazil, independent of the banana genotype, may be evidence of a fitness advantage of individuals in the *Foc* population.

The SIX genes seem to play an important role as virulence factors in *Foc* and the analysis of the profile of these genes in a population can be useful to understand race dynamics. SIX8 is required for the virulence of *Foc* in Cavendish bananas (An et al. 2019). In addition, two SIX8 homologues, SIX8a and SIX8b, are present in the genomes of race 4 strains (TR4 and SR4), but have not been reported in individuals of races 1 or 2 (Fraser-Smith et al., 2014). Specifically, SIX8a is present in all race 4 isolates (TR4 and SR4), whereas SIX8b has been reported in SR4 individuals only. The analysis of the SIX gene profile in the current study, complemented by other PCR-diagnostic methods, support the occurrence and ample distribution of subtropical race 4 among Brazilian isolates of *Foc* (An et al. 2019, Fraser-Smith et al., 2014). Overall, there was a trend of association of SIX genes profile, clades and VCGs. Given the supposedly prevalence of asexual reproduction in the evolutionary trajectory of *Foc*, some degree of association among different markers may be expected. The lack of complete association commonly seen in other asexual *F*. *oxysporum* could be the result of rare recombination events that can be mediated by compatible VCGs as discussed for the complexes herein reported.

The VCG tests could have been less laborious and more efficient if the diagnosis by PCR-RFLP for lineage classification had worked. The separation of the *Foc* strains into lineages correlates well with the vegetative compatibility group (Fourie et al. 2009). The PCR-RFLP analyses were successful in classifying the isolates into Clade A or B as proposed by Fourie et al (2009). However, the digestion that would classify the Brazilian isolates into lineages, generated atypical cleavage profiles. Karangwa et al. (2018) reported a similar result when digested some Clade A isolates with *BceA*I restriction enzyme. Although this method was consistent for the characterization of *Foc* isolates from Asia and Africa (Fourie et al. 2009; Mostert et al. 2017; Karangwa et al. 2018), it was not suitable in this study may be due to inherent genetic differences in the Brazilian population in comparison to the other strains used in the development of the method. As a consequence, we cannot safely state that strains that do not form heterokaryon with any of the VCG testers correlate to a lineage or belong to a VCG not reported yet. Previous studies have also identified strains that do not pair with testers, and the authors raised the possibility of these “un-matched” isolates to belong to novel VCGs (Mostert et al. 2017; Karangwa et al. 2018). Based on the impossibility of VCG assignment based on restriction pattern it can be speculated that there is large genetic variation in the population of *Foc* in Brazil. This was also supported by the large proportion of unknown VCGs.

All previous studies that aimed to describe the structure of the population of *Foc* in Brazil (Costa et al. 2015; Cunha et al. 2015) had limitations in relation to the amount of markers used and to the sampling scheme. These issues could have underestimated the variability in the population of the pathogen. In the present study, we attempted to minimize the effects of these limitations by expanding the sampling to other regions, including isolates from different banana producing areas such as the Ribeira Valley in São Paulo state, the Cariri Region of Ceará state, and other northern and southern regions that had not been examined. Based on the SSR genotyping, the *Foc* population appears to have low to moderate genetic variability. Clonal fraction in this study was high (77.8%), similar to that reported in another study with Brazilian isolates (74.7%) (Costa et al, 2015). MLG richness estimate for the previous study was 52 among 214 isolates and 35 among 158 isolates used in the present study. Similarly, the genotypic diversity estimated by the Stoddart & Taylor index in the studies was 5.96 and 3.96, for the study conducted by Costa et al (2015) and the present one, respectively. Even though sample sizes differ, estimates are anticipated to be similar if a rarefaction analysis is conducted. Overall, these results suggest that *Foc* fits a model of predominant clonal evolution (PCE). The assumptions of the PCE model are strong linkage disequilibrium (LD), widespread occurrence of stable MLG, and restrained recombination but not absent (Tibayrenc and Ayala 2017, 2021).

Finally, there is an urgent need to develop a larger set of SSR markers for *Foc* or, preferentially, use genome wide snps to increase the resolution of the population genetics analyses. A crucial question that would benefit from higher-resolution marker datasets is how clonal is the population of *Foc*? Is there evidence for the persistence of lineages with mutants at few loci or is there evidence for a more variable population of Foc due to genetic exchange occurring more often than expected? These are important questions related to the durability of resistant varieties, the main disease control measure against FWB worldwide.

## Acknowledgments

This research was supported by the Fundação de Amparo à Pesquisa do Estado de Minas Gerais - FAPEMIG. We thank Dr. Randy Ploetz and Joshua Konkol for supplying strains of known VCG testers of *Fusarium oxysporum* f. sp. *cubense;* and Evelyn L. Coca for her contribution to the execution in part of the VCG assays.

