## Supplemental table 1, 2, 3 and 4 for "The Brazilian population of *Fusarium oxysporum* f. sp. *cubense* is not structured by VCG or by geographic origin"

**Supplementary table 1** Isolates of *Fusarium oxysporum* f. sp. *cubense.*

| **Isolate**  **code** | **Year** | **Collector/**  **Sponsor** | **Municipality** | **Region** | **State** | **Longitude** | **Latitude** | **Altitude** | **Host** | **Cultivar** | **Genotype** | **Clado** | **VCG** | **SSR Genotyping** | **SIX profile** |
| --- | --- | --- | --- | --- | --- | --- | --- | --- | --- | --- | --- | --- | --- | --- | --- |
| UFV-Foc001 | 2015 | Miguel Dita | Jacupiranga | Southeast | SP | -48,0957780 | -24,8858060 | 33 | Banana | Gran Naine | AAA | A | 0120 | + | 1, 8, 8a and 8b |
| UFV-Foc002 | 2015 | Miguel Dita | Jacupiranga | Southeast | SP | -48,0551330 | -24,8860330 | 33 | Banana | Gallil 7 | AAA | A | 0120 | - | 1,7,8, 8a and 8b |
| UFV-Foc003 | 2016 | Miguel Dita | Cajati | Southeast | SP | -48,2214720 | -24,7013780 | 168 | Banana | Prata anã | AAA | A | 0120 | + | 1,7,8, 8a and 8b |
| UFV-Foc013 | 2016 | Miguel Dita | Jacupiranga | Southeast | SP | -48,0551330 | -24,8860330 | 148 | Banana | Prata comum | AAB | A | 0120 | - | 1,7,8, 8a and 8b |
| UFV-Foc016 | 2016 | Miguel Dita | Jacupiranga | Southeast | SP | -48,0551330 | -24,8860330 | 148 | Banana | Prata comum | AAB | A | 0120/15 | + | 1,7,8, 8a and 8b |
| UFV-Foc024 | 2016 | Miguel Dita | Pariquera-Açú | Southeast | SP | -47,8853080 | -24,6181440 | 47 | Banana | Prata Catarina | AAB | A | 0120 | + | 1,7,8, 8a and 8b |
| UFV-Foc029 | 2016 | Miguel Dita | Pariquera-Açú | Southeast | SP | -47,8853080 | -24,6181440 | 47 | Banana | Prata Catarina | AAB | A | 0120 | - | 1,7,8, 8a and 8b |
| UFV-Foc032 | 2016 | Miguel Dita | Eldorado | Southeast | SP | -48,2023830 | -24,6296140 | 92 | Banana | Prata Anã | AAB | A | 0120 | - | 1,7,8, 8a and 8b |
| UFV-Foc033 | 2016 | Miguel Dita | Eldorado | Southeast | SP | -48,2023830 | -24,6296140 | 92 | Banana | Prata Anã | AAB | A | 0120 | - | 1,7,8, 8a and 8b |
| UFV-Foc047 | 2016 | Miguel Dita | Jacupiranga | Southeast | SP | -48,0551330 | -24,8860330 | 33 | Banana | Gallil 7 | AAA | A | 0120 | - | 1,7,8, 8a and 8b |
| UFV-Foc053 | 2016 | Miguel Dita | Marinópolis | Southeast | SP | -50,8332250 | -20,4611890 | 418 | Banana | Maçã | AAB | A | Unknown | + | 1 and 13 |
| UFV-Foc054 | 2016 | Miguel Dita | Marinópolis | Southeast | SP | -50,8332250 | -20,4611890 | 418 | Banana | Maçã | AAB | A | 0120 | - | - |
| UFV-Foc059 | 2016 | Miguel Dita | Marinópolis | Southeast | SP | -50,8333110 | -20,4611830 | 418 | Banana | Maçã | AAB | A | Unknown | + | 1 and 13 |
| UFV-Foc061 | 2016 | Miguel Dita | Marinópolis | Southeast | SP | -50,8167080 | -20,4366500 | 389 | Banana | Maçã | AAB | A | Unknown | + | 1, 8, 8a and 8b |
| UFV-Foc063 | 2016 | Miguel Dita | Marinópolis | Southeast | SP | -50,8166030 | -20,4365940 | 389 | Banana | Maçã | AAB | A | Unknown | + | 1 and 13 |
| UFV-Foc069 | 2016 | Miguel Dita | Marinópolis | Southeast | SP | -50,8258610 | -20,4348440 | 406 | Banana | Maçã | AAB | A | Unknown | + | 1 and 13 |
| UFV-Foc072 | 2016 | Miguel Dita | Marinópolis | Southeast | SP | -50,8258080 | -20,4348830 | 406 | Banana | Maçã | AAB | A | Unknown | + | 1,7,8, 8a and 8b |
| UFV-Foc078 | 2016 | Miguel Dita | Marinópolis | Southeast | SP | -50,8042920 | -20,4398170 | 400 | Banana | Maçã | AAB | A | Unknown | + | No six detected |
| UFV-Foc081 | 2016 | Miguel Dita | Marinópolis | Southeast | SP | -50,8040420 | -20,4397170 | 400 | Banana | Maçã | AAB | A | *crn* mutant | *+* | 1 and 13 |
| UFV-Foc085 | 2016 | Miguel Dita | Marinópolis | Southeast | SP | -50,8332250 | -20,4611890 | 418 | Banana | Maçã | AAB | A | 0120 | - | - |
| UFV-Foc086 | 2016 | Miguel Dita | Aparecida D´Oeste | Southeast | SP | -50,8767420 | -20,4798470 | 370 | Banana | Maçã | AAB | A | NT | + | 1,7,8, 8a, 8b and 13 |
| UFV-Foc087 | 2016 | Miguel Dita | Corupá | South | SC | -49,2433330 | -26,4261110 | 52 | Banana | Prata | AAB | A | Unknown | + | 1,7,8, 8a, 8b and 13 |
| UFV-Foc088 | 2016 | Miguel Dita | Corupá | South | SC | -49,2433330 | -26,4261110 | 52 | Banana | Nanica | AAA | A | 0120 | + | 1,7,8, 8a and 8b |
| UFV-Foc098 | 2016 | Miguel Dita | São Bento do Sapucaí | Southeast | SP | -45,7007690 | -22,6884440 | 1045 | Banana | Prata | AAB | A | 0120 | + | 1,7,8, 8a and 8b |
| UFV-Foc099 | 2016 | Miguel Dita | São Bento do Sapucaí | Southeast | SP | -45,7007690 | -22,6884440 | 1045 | Banana | Prata | AAB | A | 0120 | - | - |
| UFV-Foc100 | 2016 | Miguel Dita | São Bento do Sapucaí | Southeast | SP | -45,7007690 | -22,6884440 | 1045 | Banana | Prata | AAB | A | 0120 | + | 1 and 2 |
| UFV-Foc101 | 2016 | Miguel Dita | São Bento do Sapucaí | Southeast | SP | -45,7007690 | -22,6884440 | 1045 | Banana | Prata Mineira | AAB | A | 0120/15 | + | 1,7,8, 8a and 8b |
| UFV-Foc111 | 2016 | Miguel Dita | São Bento do Sapucaí | Southeast | SP | -45,6968170 | -22,6873970 | 1111 | Banana | Prata Mineira | AAB | A | 0120 | + | 1,7,8, 8a, 8b and 13 |
| UFV-Foc113 | 2016 | Miguel Dita | São Bento do Sapucaí | Southeast | SP | -45,6967140 | -22,6874170 | 1114 | Banana | Prata Mineira | AAB | A | 0120/15 | + | 1,7,8, 8a, 8b and 13 |
| UFV-Foc114 | 2016 | Miguel Dita | São Bento do Sapucaí | Southeast | SP | -45,6967140 | -22,6874170 | 1114 | Banana | Prata Mineira | AAB | A | 0120/15 | + | 1,7,8, 8a and 8b |
| UFV-Foc117 | 2016 | Miguel Dita | São Bento do Sapucaí | Southeast | SP | -45,6967140 | -22,6874170 | 1114 | Banana | Prata Mineira | AAB | A | 0120/15 | + | 1,7,8, 8a, 8b and 13 |
| UFV-Foc122 | 2016 | Miguel Dita | São Bento do Sapucaí | Southeast | SP | -45,6967140 | -22,6874170 | 1114 | Banana | Prata Mineira | AAB | A | 0120 | - | 1,7,8, 8a and 8b |
| UFV-Foc123 | 2016 | Miguel Dita | São Bento do Sapucaí | Southeast | SP | -45,6967140 | -22,6874170 | 1114 | Banana | Prata Mineira | AAB | A | 0120 | + | 1,7,8, 8a and 8b |
| UFV-Foc124 | 2016 | Miguel Dita | São Bento do Sapucaí | Southeast | SP | -45,6967140 | -22,6874170 | 1114 | Banana | Prata Mineira | AAB | A | 0120 | + | 1,7,8, 8a and 8b |
| UFV-Foc125 | 2016 | Miguel Dita | São Bento do Sapucaí | Southeast | SP | -45,6967140 | -22,6874170 | 1114 | Banana | Prata Mineira | AAB | A | 0120 | - | 1,7,8, 8a and 8b |
| UFV-Foc130 | 2016 | Miguel Dita | São Bento do Sapucaí | Southeast | SP | -45,6967140 | -22,6874170 | 1114 | Banana | Prata Mineira | AAB | A | 0120 | - | - |
| UFV-Foc136 | 2016 | Miguel Dita | Santa Mariana | South | PR | -50,5186360 | -23,1838650 | 507 | Banana | Maçã | AAB | A | Unknown | + | 1 and 13 |
| UFV-Foc142 | 2016 | Miguel Dita | Santa Mariana | South | PR | -50,5186360 | -23,1838650 | 507 | Banana | Maçã | AAB | A | Unknown | + | 1 and 13 |
| UFV-Foc144 | 2016 | Miguel Dita | Santa Mariana | South | PR | -50,5465310 | -22,9533090 | 411 | Banana | Maçã | AAB | A | Unknown | + | 1 and 13 |
| UFV-Foc145 | 2016 | Miguel Dita | Santa Mariana | South | PR | -50,5465310 | -22,9533090 | 411 | Banana | Maçã | AAB | - | NT | + | 1 and 13 |
| UFV-Foc146 | 2016 | Miguel Dita | Santa Mariana | South | PR | -50,5465310 | -22,9533090 | 411 | Banana | Maçã | AAB | - | NT | + | 1 and 13 |
| UFV-Foc151 | 2015 | Miguel Dita | Cândido Mota | Southeast | SP | -50,4834570 | -22,7365170 | 465 | Banana | Maçã | AAB | A | NT | + | 1 and 13 |
| UFV-Foc152 | - | Miguel Dita | Penápolis | Southeast | SP | -49,6717810 | -21,2433780 | 383 | Banana | Maçã | AAB | A | Unknown | + | 1 and 13 |
| UFV-Foc154 | - | Miguel Dita | Penápolis | Southeast | SP | -50,1089860 | -21,2272530 | 383 | Banana | Maçã | AAB | A | Unknown | + | 1 and 13 |
| UFV-Foc161 | 2016 | Miguel Dita | Penápolis | Southeast | SP | -50,1089860 | -21,2272530 | 383 | Banana | Maçã | AAB | A | Unknown | + | 1 and 13 |
| UFV-Foc166 | 2016 | Miguel Dita | Penápolis | Southeast | SP | -50,1087810 | -21,2272860 | 381 | Banana | Maçã | AAB | A | Unknown | + | 1 and 13 |
| UFV-Foc171 | 2016 | Miguel Dita | Penápolis | Southeast | SP | -50,1085220 | -21,2270000 | 380 | Banana | Maçã | AAB | A | *crn* mutant | *-* | 1 and 13 |
| UFV-Foc182 | 2016 | Miguel Dita | Penápolis | Southeast | SP | -49,6717810 | -21,2433780 | 420 | Banana | Maçã | AAB | A | Unknown | + | 1 and 13 |
| UFV-Foc191 | 2016 | Miguel Dita | Penápolis | Southeast | SP | -49,6717810 | -21,2444890 | 413 | Banana | Maçã | AAB | A | Unknown | + | 1 and 13 |
| UFV-Foc201 | 2016 | Miguel Dita | Penápolis | Southeast | SP | -49,4732920 | -21,4149860 | 421 | Banana | Maçã | AAB | A | Unknown | + | 1 and 13 |
| UFV-Foc210 | 2016 | Daniel Heck/ Eduardo Mizubuti | São Bento do Sapucaí | Southeast | SP | -45,7363890 | -22,6883330 | 874 | Banana | Prata | AAB | A | 0120 | + | 1, 8a and 8b |
| UFV-Foc212 | 2016 | Daniel Heck/ Eduardo Mizubuti | Penápolis | Southeast | SP | -50,0780560 | -21,4208330 | 407 | Banana | Maçã | AAB | A | Unknown | + | 1 and 13 |
| UFV-Foc213 | 2016 | Daniel Heck/ Eduardo Mizubuti | Santa Mariana | South | PR | -50,4880560 | -23,1413890 | 421 | Banana | Maça | AAB | A | 0120 | + | 13 |
| UFV-Foc215 | 2016 | Daniel Heck/ Eduardo Mizubuti | Jacupiranga | Southeast | SP | -48,0988770 | -24,8902540 | 102 | Banana | Prata | AAB | A | 0120 | + | 1,7,8, 8a, 8b and 13 |
| UFV-Foc216 | 2016 | Daniel Heck/ Eduardo Mizubuti | Jacupiranga | Southeast | SP | -48,0988770 | -24,8902540 | 102 | Banana | Nanica | AAA | A | 0120/15 | + | 1,7,8, 8a and 8b |
| UFV-Foc217 | - | Fernando Haddad | Cruz das Almas | Northeast | BA | -39,1219440 | -12,6530560 | 212 | Banana | Maçã | AAB | B | Unknown | + | 1, 8a, 8b and 13 |
| UFV-Foc218 | - | Fernando Haddad | Itajuípi | Northeast | BA | -39,3770000 | -14,6747000 | 108 | Banana | Prata-Anã | AAB | - | NT | + | 1,7,8, 8a, 8b and 13 |
| UFV-Foc223 | 2011 | Fernando Haddad | Porteirinha | Southeast | MG | -43,3733300 | -15,9233330 | 556 | Banana | Prata anã | AAB | - | NT | - | 1 and 13 |
| UFV-Foc227 | 2011 | Fernando Haddad | Guanambi | Northeast | BA | -42,8736100 | -14,4652780 | 515 | Banana | Prata anã | AAB | - | NT | - | 1,7,8, 8a and 8b |
| UFV-Foc242 | 2011 | Fernando Haddad | Tancredo Neves | Northeast | BA | -39,5191700 | -13,5822220 | 253 | Banana | Maçã | AAB | - | NT | + | 1 and 13 |
| UFV-Foc243 | 2011 | Fernando Haddad | Tancredo Neves | Northeast | BA | -39,5230600 | -13,5722220 | 253 | Banana | Maçã | AAB | - | NT | - | 1, 8b and 13 |
| UFV-Foc244 | 2011 | Fernando Haddad | Porteirinha | Southeast | MG | -43,3733300 | -15,9233330 | 556 | Banana | Prata anã | AAB | - | NT | + | 1 and 13 |
| UFV-Foc245 | 2011 | Fernando Haddad | Juazeiro | Northeast | BA | -40,3991700 | -9,4694440 | 369 | Banana | Maçã | AAB | - | NT | + | 1, 8b and 13 |
| UFV-Foc248 | 2011 | Fernando Haddad | Juazeiro | Northeast | BA | -40,3411100 | -9,5447220 | 369 | Banana | Maçã | AAB | - | NT | + | 1 and 13 |
| UFV-Foc249 | 2011 | Fernando Haddad | Juazeiro | Northeast | BA | -40,4261100 | -9,5144440 | 369 | Banana | Maçã | AAB | - | NT | + | 1, 8b and 13 |
| UFV-Foc250 | 2011 | Fernando Haddad | Ponto Novo | Northeast | BA | -40,3663900 | -10,9502780 | 368 | Banana | Prata anã | AAB | - | NT | + | 1, 8b and 13 |
| UFV-Foc251 | 2011 | Fernando Haddad | Lavras | Southeast | MG | -44,9913900 | -21,2313890 | 919 | Banana | Maçã | AAB | - | NT | + | 1, 8b and 13 |
| UFV-Foc252 | 2011 | Fernando Haddad | Lavras | Southeast | MG | -44,9913900 | -21,2313890 | 919 | Banana | Maçã | AAB | - | NT | + | 1 |
| UFV-Foc256 | 2011 | Fernando Haddad | Guanambi | Northeast | BA | -42,8663900 | -14,8647220 | 515 | Banana | Prata Anã | AAB | - | NT | + | 1,7, 8b and 13 |
| UFV-Foc262 | 2011 | Fernando Haddad | Itajuípe | Northeast | BA | -39,3834100 | -14,9663890 | 108 | Banana | Maçã | AAB | - | NT | + | 1,7, 8b and 13 |
| UFV-Foc264 | 2011 | Fernando Haddad | Itajuípe | Northeast | BA | -39,4269400 | -14,7630560 | 108 | Banana | Prata | AAB | - | NT | - | 1, 8b and 13 |
| UFV-Foc265 | 2011 | Fernando Haddad | Itajuípe | Northeast | BA | -39,4241700 | -14,7566670 | 108 | Banana | Prata | AAB | - | NT | + | 1, 8b and 13 |
| UFV-Foc266 | 2011 | Fernando Haddad | Itajuípe | Northeast | BA | -39,6180600 | -14,7708330 | 108 | Banana | Prata | AAB | - | NT | + | 1, 7, 8b and 13 |
| UFV-Foc267 | 2011 | Fernando Haddad | Gandu | Northeast | BA | -39,7100000 | -13,9477780 | 155 | Banana | Prata | AAB | - | NT | + | 1, 7, 8b and 13 |
| UFV-Foc268 | 2011 | Fernando Haddad | Wenceslau | Northeast | BA | -39,9086100 | -13,7922220 | 146 | Banana | Prata | AAB | - | NT | - | 1, 7 and 8b |
| UFV-Foc269 | 2011 | Fernando Haddad | Teolândia | Northeast | BA | -39,5841700 | -13,7911110 | 209 | Banana | Prata | AAB | - | NT | + | 1, 8b and 13 |
| UFV-Foc270 | 2011 | Fernando Haddad | Pedra Branca | Northeast | CE | -39,7166700 | -5,4508330 | 500 | Banana | Maçã | AAB | A | Unknown | + | 1 and 13 |
| UFV-Foc271 | 2011 | Fernando Haddad | Mossoró | Northeast | RN | -37,3438900 | -5,1877780 | 500 | Banana | Tropical | AAAB | - | Unknown | - | 1 and 13 |
| UFV-Foc272 | 2011 | Fernando Haddad | Dom Pedro de Alcântara | South | RS | -49,8716400 | -29,3524440 | 7 | Banana | Prata | AAB | - | Unknown | + | 1, 7 and 8 |
| UFV-Foc273 | 2011 | Fernando Haddad | Mâmpituba | South | RS | -49,9354400 | -29,3960560 | 37 | Banana | Prata | AAB | - | NT | + | 1, 7 and 8 |
| UFV-Foc274 | 2011 | Fernando Haddad | Três Cachoeiras | South | RS | -49,9244400 | -29,4555560 | 9 | Banana | Prata | AAB | - | NT | + | - |
| UFV-Foc275 | 2011 | Fernando Haddad | Três Cachoeiras | South | RS | -49,9244400 | -29,4555560 | 39 | Banana | Prata | AAB | A | 0120/15 | + | 1, 8 and 8b |
| UFV-Foc276 | 2011 | Fernando Haddad | Ponto Novo | Northeast | BA | -40,3663900 | -10,9502780 | 368 | Banana | Prata Gorutuba | AAB | - | NT | + | 1, 7 and 13 |
| UFV-Foc277 | 2011 | Fernando Haddad | Ponto Novo | Northeast | BA | -40,3663900 | -10,9502780 | 368 | Banana | Thap Maeo | AAB | - | NT | + | 1, 7, 8b and 13 |
| UFV-Foc278 | 2011 | Fernando Haddad | Ponto Novo | Northeast | BA | -40,3663900 | -10,9502780 | 368 | Banana | Prata Gorutuba | AAB | - | NT | + | 1, 7, 8b and 13 |
| UFV-Foc297 | 2014 | Fernando Haddad | Eldorado | Southeast | SP | -48,2023830 | -24,6296140 | 92 | Banana | Nanica | AAA | A | 0120 | + | 1, 7 and 8 |
| UFV-Foc298 | 2014 | Fernando Haddad | Pedro de Toledo | Southeast | SP | -47,2336000 | -24,2768000 | 44 | Banana | Nanica | AAA | B | 0124/22 | + | 1 and 13 |
| UFV-Foc303 | 2014 | Fernando Haddad | Miracatu | Southeast | SP | -47,4585000 | -24,2860000 | 27 | Banana | Prata-Anã | AAB | A | 0120 | + | 1,7,8, 8a and 8b |
| UFV-Foc305 | 2014 | Fernando Haddad | Pedro de Toledo | Southeast | SP | -47,2336000 | -24,2768000 | 44 | Banana | Prata-Anã | AAB | A | 0120 | + | 1,7,8, 8a and 8b |
| UFV-Foc309 | 2014 | Fernando Haddad | Jacupiranga | Southeast | SP | -48,0988770 | -24,8902540 | 33 | Banana | Nanica | AAA | A | 0120 | + | 1,7,8, 8a and 8b |
| UFV-Foc319 | 2014 | Fernando Haddad | Pariquera-Açu | Southeast | SP | -48,0988770 | -24,8902540 | 33 | Banana | Prata Catarina | AAB | A | 0120 | - | 1,7,8, 8a and 8b |
| UFV-Foc332 | 2014 | Fernando Haddad | Sete Barras | Southeast | SP | -47,9280000 | -24,3845000 | 30 | Banana | Nanica | AAA | A | 0120 | - | 1,7,8, 8a and 8b |
| UFV-Foc334 | 2014 | Fernando Haddad | Palmitos | South | SC | -53,0929000 | -27,0402000 | 406 | Banana | Nanicão | AAA | - | NT | + | - |
| UFV-Foc342 | 2014 | Fernando Haddad | Registro | Southeast | SP | -47,8494000 | -24,5083000 | 25 | Banana | Prata Comum | AAB | A | 0120 | + | 1, 7, 8, 8a and 8b |
| UFV-Foc354 | 2014 | Fernando Haddad | Corupá | South | SC | -49,2430560 | -26,4252780 | 75 | Banana | Nanica | AAA | A | 0120 | + | 1 and 13 |
| UFV-Foc355 | 2014 | Fernando Haddad | Corupá | South | SC | -49,2826270 | -26,4567230 | 52 | Banana | Prata-Anã | AAB | A | *crn* mutant | *+* | *-* |
| UFV-Foc356 | 2014 | Fernando Haddad | Corupá | South | SC | -49,2826270 | -26,4567230 | 52 | Banana | Prata-Anã | AAB | A | Unknown | + | 1, 8b and 13 |
| UFV-Foc358 | 2014 | Fernando Haddad | Corupá | South | SC | -49,2826270 | -26,4567230 | 52 | Banana | Nanica | AAA | A | 0120 | + | 1, 8a and 8b |
| UFV-Foc387 | - | Fernando Haddad | Bom Jesus da Lapa | Northeast | BA | -43,2505000 | -13,1518000 | 436 | Banana | Prata Gorotuba | AAB | - | NT | + | - |
| UFV-Foc412 | 2008 | Sami Michereff | Bom Conselho | Northeast | PE | -36,6833330 | -9,1641670 | 630 | Banana | - | - | B | 0124 | + | 1, 12 and 13 |
| UFV-Foc413 | 2006 | Sami Michereff | Vicencia | Northeast | PE | -35,3275000 | -7,6683330 | 157 | Banana | - | - | A | Unknown | + | 1, 7, 8, 12 and 13 |
| UFV-Foc438 | 2006 | Sami Michereff | Cruz das Almas | Northeast | BA | -39,1219440 | -12,6530560 | 212 | Banana | - | - | A | 0120 | + | 1, 12 and 13 |
| UFV-Foc440 | 2004 | Sami Michereff | Campina Grande | Northeast | PB | -35,8816670 | -7,2308330 | 512 | Algodão | - | - | - | NT | + | - |
| UFV-Foc470 | 2007 | Sami Michereff | Cruz das Almas | Northeast | BA | -39,1219440 | -12,6530560 | 212 | Banana | - | - | - | Unknown | + | 1 and 13 |
| UFV-Foc471 | 2007 | Sami Michereff | Cruz das Almas | Northeast | BA | -39,1219440 | -12,6530560 | 212 | Banana | - | - | - | Unknown | + | 1 and 13 |
| UFV-Foc473 | 2006 | Sami Michereff | Rio Largo | Northeast | AL | -35,8580560 | -9,4802780 | 44 | Banana | - | - | - | Unknown | + | 1 and 13 |
| UFV-Foc478 | 2006 | Sami Michereff | Rio Largo | Northeast | AL | -35,8580560 | -9,4802780 | 44 | Banana | - | - | - | Unknown | + | 1 and 13 |
| UFV-Foc480 | 2006 | Sami Michereff | Rio Largo | Northeast | AL | -35,8580560 | -9,4802780 | 44 | Banana | - | - | - | Unknown | + | 1 and 13 |
| UFV-Foc486 | 2010 | Sami Michereff | Recife | Northeast | PE | -34,8813890 | -8,0541670 | 7 | Crotalária | - | - | - | NT | + | 1 and 13 |
| UFV-Foc511 | 2017 | Daniel Heck/ Eduardo Mizubuti | Janaúba | Southeast | MG | -43,3521700 | -15,7644800 | 524 | Banana | Prata Anã | AAB | A | 0120 | + | 1, 7, 8, 8a and 8b |
| UFV-Foc512 | 2017 | Daniel Heck/ Eduardo Mizubuti | Jaíba | Southeast | MG | -43,7885320 | -15,1791840 | 475 | Banana | Prata Catarina | AAB | A | 0120 | + | 1, 7, 8, 8a and 8b |
| UFV-Foc513 | 2017 | Daniel Heck/ Eduardo Mizubuti | Jaíba | Southeast | MG | -43,7885320 | -15,1791840 | 475 | Banana | Prata Catarina | AAB | A | 0120 | + | 1, 7, 8, 8a and 8b |
| UFV-Foc514 | 2017 | Daniel Heck/ Eduardo Mizubuti | Jaíba | Southeast | MG | -43,7880210 | -15,1766520 | 474 | Banana | Prata Gorutuba | AAB | A | 0120 | + | 1, 7, 8, 8a and 8b |
| UFV-Foc515 | 2017 | Daniel Heck/ Eduardo Mizubuti | Jaíba | Southeast | MG | -43,7873160 | -15,1748270 | 473 | Banana | Prata Catarina | AAB | A | 0120 | + | 1, 8a and 8b |
| UFV-Foc516 | 2017 | Daniel Heck/ Eduardo Mizubuti | Serra do Ramalho | Northeast | BA | -43,7021950 | -13,2359800 | 449 | Banana | Prata Gorutuba | AAB | A | *crn* mutant | *+* | 1, 7, 8, 8a and 8b |
| UFV-Foc517 | 2017 | Daniel Heck/ Eduardo Mizubuti | Serra do Ramalho | Northeast | BA | -43,7021950 | -13,2359800 | 449 | Banana | Prata Gorutuba | AAB | A | 0120 | + | 1, 7, 8, 8a and 8b |
| UFV-Foc518 | 2017 | Daniel Heck/ Eduardo Mizubuti | Serra do Ramalho | Northeast | BA | -43,7021950 | -13,2359800 | 449 | Banana | Prata Gorutuba | AAB | A | 0120 | + | 1, 7, 8, 8a and 8b |
| UFV-Foc519 | 2017 | Daniel Heck/ Eduardo Mizubuti | Janaúba | Southeast | MG | -43,3503930 | -15,7668810 | 528 | Banana | Prata Catarina | AAB | A | 0120 | + | 1, 7, 8, 8a and 8b |
| UFV-Foc520 | 2017 | Daniel Heck/ Eduardo Mizubuti | Jaíba | Southeast | MG | -43,7880210 | -15,1766520 | 474 | Banana | Prata Gorutuba | AAB | - | NT | + | 1, 7, 8, 8a and 8b |
| UFV-Foc521 | 2017 | Daniel Heck/ Eduardo Mizubuti | Janaúba | Southeast | MG | -43,3521700 | -15,7644800 | 524 | Banana | Prata Anã | AAB | - | NT | + | 1, 7, 8, 8a and 8b |
| UFV-Foc522 | 2017 | Daniel Heck/ Eduardo Mizubuti | Jaíba | Southeast | MG | -43,7885320 | -15,1791840 | 475 | Banana | Prata Catarina | AAB | - | NT | + | 1, 7, 8, 8a and 8b |
| UFV-Foc523 | 2017 | Daniel Heck/ Eduardo Mizubuti | Jaíba | Southeast | MG | -43,7880210 | -15,1766520 | 474 | Banana | Prata Gorutuba | AAB | - | NT | + | 1, 8, 8a and 8b |
| UFV-Foc526 | 2017 | Daniel Heck/ Eduardo Mizubuti | Corupá | South | SC | -49,2927850 | -26,4567230 | 140 | Banana | Caturra | AAA | A | 0120 | + | 1, 7, 8, 8a and 8b |
| UFV-Foc529 | 2017 | Daniel Heck/ Eduardo Mizubuti | Corupá | South | SC | -49,2927850 | -26,4567230 | 140 | Banana | Caturra | AAA | A | 0120 | + | 1, 7, 8, 8a and 8b |
| UFV-Foc530 | 2017 | Daniel Heck/ Eduardo Mizubuti | Corupá | South | SC | -49,2826270 | -26,4413000 | 113 | Banana | Nanicão | AAA | A | 0120 | + | 1, 7, 8, 8a and 8b |
| UFV-Foc531 | 2017 | Daniel Heck/ Eduardo Mizubuti | Corupá | South | SC | -49,2826270 | -26,4413000 | 113 | Banana | Nanicão | AAA | - | NT | + | 1, 7, 8, 8a and 8b |
| UFV-Foc533 | 2017 | Daniel Heck/ Eduardo Mizubuti | Corupá | South | SC | -49,2826270 | -26,4413000 | 113 | Banana | Nanicão | AAA | - | NT | + | 1, 7, 8, 8a and 8b |
| UFV-Foc534 | 2017 | Daniel Heck/ Eduardo Mizubuti | Corupá | South | SC | -49,2985920 | -26,4772360 | 166 | Banana | Prata Anã | AAB | A | 0120 | + | 1, 7, 8, 8a and 8b |
| UFV-Foc535 | 2017 | Daniel Heck/ Eduardo Mizubuti | Corupá | South | SC | -49,2985920 | -26,4413000 | 166 | Banana | Prata Anã | AAB | - | NT | + | 1, 8, 8a and 8b |
| UFV-Foc536 | 2017 | Daniel Heck/ Eduardo Mizubuti | Jaraguá do Sul | South | SC | -49,1896930 | -26,4378210 | 45 | Banana | Prata Catarina | AAB | A | 0120/15 | + | 1, 8, 8a and 8b |
| UFV-Foc537 | 2017 | Daniel Heck/ Eduardo Mizubuti | Jaraguá do Sul | South | SC | -49,1896930 | -26,4378210 | 45 | Banana | Prata Catarina | AAB | - | NT | + | 1 |
| UFV-Foc539 | 2017 | Daniel Heck/ Eduardo Mizubuti | Corupá | South | SC | -49,2430560 | -26,4252780 | 75 | Banana | Figo | ABB | A | 0120/15 | + | 1, 7, 8, 8a and 8b |
| UFV-Foc540 | 2017 | Daniel Heck/ Eduardo Mizubuti | Serra do Ramalho | Northeast | BA | -43,7021950 | -13,2359800 | 449 | Banana | Prata Gorutuba | AAB | - | NT | + | - |
| UFV-Foc541 | 2017 | Daniel Heck/ Eduardo Mizubuti | Serra do Ramalho | Northeast | BA | -43,7021950 | -13,2359800 | 449 | Banana | Prata Gorutuba | AAB | - | Unknown | + | 1, 8a and 8b |
| UFV-Foc542 | 2017 | Daniel Heck/ Eduardo Mizubuti | Janaúba | Southeast | MG | -43,3521700 | -15,7644800 | 524 | Banana | Prata Anã | AAB | B | *crn* mutant | *+* | 1, 7, 8, 8a and 8b |
| UFV-Foc543 | 2017 | Daniel Heck/ Eduardo Mizubuti | Jaíba | Southeast | MG | -43,7873160 | -15,1748270 | 473 | Banana | Prata Catarina | AAB | B | 0124 | + | 1 |
| UFV-Foc557 | 2017 | Daniel Heck/ Eduardo Mizubuti | Janaúba | Southeast | MG | -43,3521700 | -15,7644800 | 524 | Banana | Prata Anã | AAB | - | NT | + | - |
| UFV-Foc559 | 2017 | Daniel Heck/ Eduardo Mizubuti | Janaúba | Southeast | MG | -43,3503930 | -15,7668810 | 528 | Banana | Prata Catarina | AAB | - | NT | + | 1, 7, 8, 8a and 8b |
| UFV-Foc560 | 2017 | Daniel Heck/ Eduardo Mizubuti | Janaúba | Southeast | MG | -43,3503930 | -15,7668810 | 528 | Banana | Prata Catarina | AAB | - | NT | + | 1, 8, 8a and 8b |
| UFV-Foc561 | 2017 | Daniel Heck/ Eduardo Mizubuti | Jaíba | Southeast | MG | -43,7880210 | -15,1766520 | 474 | Banana | Prata Gorutuba | AAB | - | NT | + | 1 and 7 |
| UFV-Foc562 | 2017 | Daniel Heck/ Eduardo Mizubuti | Corupá | South | SC | -49,2985920 | -26,4772360 | 166 | Banana | Prata Anã | AAB | - | NT | + | 1, 8, 8a and 8b |
| UFV-Foc569 | 2017 | Daniel Heck/ Eduardo Mizubuti | Corupá | South | SC | -49,2826270 | -26,4413000 | 113 | Banana | Nanicão | AAA | - | NT | + | 1, 7 and 8b |
| UFV-Foc571 | 2017 | Daniel Heck/ Eduardo Mizubuti | Corupá | South | SC | -49,1896930 | -26,4378210 | 45 | Banana | Prata Catarina | AAB | - | NT | + | 1, 8, 8a and 8b |
| UFV-Foc572 | 2017 | Daniel Heck/ Eduardo Mizubuti | Janaúba | Southeast | MG | -43,3521700 | -15,7644800 | 524 | Banana | Prata Anã | AAB | - | NT | + | 1, 7 and 8b |
| UFV-Foc575 | 2017 | Daniel Heck/ Eduardo Mizubuti | Jaíba | Southeast | MG | -43,7880210 | -15,1766520 | 474 | Banana | Prata Gorutuba | AAB | - | NT | + | 1, 7 and 8b |
| UFV-Foc576 | 2017 | Daniel Heck/ Eduardo Mizubuti | Jaíba | Southeast | MG | -43,7873160 | -15,1748270 | 473 | Banana | Prata Catarina | AAB | A | 0120 | + | 1, 8a and 8b |
| UFV-Foc577 | 2017 | Daniel Heck/ Eduardo Mizubuti | Teixeiras | Southeast | MG | -42,8337730 | -20,6384300 | 690 | Banana | Prata | AAB | A | 0120 | + | 1, 8a and 8b |
| UFV-Foc578 | 2017 | Daniel Heck/ Eduardo Mizubuti | Teixeiras | Southeast | MG | -42,8337730 | -20,6384300 | 690 | Banana | Prata | AAB | A | 0120 | + | 1, 8, 8a and 8b |
| UFV-Foc579 | 2017 | Daniel Heck/ Eduardo Mizubuti | Teixeiras | Southeast | MG | -42,8337730 | -20,6384300 | 690 | Banana | Prata | AAB | B | *crn* mutant | *+* | 1, 7, 8, 8a and 8b |
| UFV-Foc580 | 2017 | Daniel Heck/ Eduardo Mizubuti | Teixeiras | Southeast | MG | -42,8337730 | -20,6384300 | 690 | Banana | Prata | AAB | - | NT | + | 7 |
| UFV-Foc581 | 2017 | Daniel Heck/ Eduardo Mizubuti | Teixeiras | Southeast | MG | -42,8337730 | -20,6384300 | 690 | Banana | Prata | AAB | A | 01215 | + | 1, 7, 8, 8a, 8b and 13 |
| UFV-Foc582 | 2017 | Daniel Heck/ Eduardo Mizubuti | Teixeiras | Southeast | MG | -42,8337730 | -20,6384300 | 690 | Banana | Prata | AAB | A | 0120 | + | 1, 8, 8a and 8b |
| UFV-Foc583 | 2017 | Daniel Heck/ Eduardo Mizubuti | Teixeiras | Southeast | MG | -42,8337730 | -20,6384300 | 690 | Banana | Prata | AAB | A | 0120 | + | 1, 8a and 8b |
| UFV-Foc584 | 2017 | Daniel Heck/ Eduardo Mizubuti | Teixeiras | Southeast | MG | -42,8337730 | -20,6384300 | 690 | Banana | Prata | AAB | - | NT | + | 1, 7 and 8b |
| UFV-Foc585 | 2017 | Daniel Heck/ Eduardo Mizubuti | Teixeiras | Southeast | MG | -42,8337730 | -20,6384300 | 690 | Banana | Prata | AAB | - | NT | + | 1 and 7 |
| UFV-Foc586 | 2017 | Daniel Heck/ Eduardo Mizubuti | Teixeiras | Southeast | MG | -42,8337730 | -20,6384300 | 690 | Banana | Prata | AAB | - | NT | + | 1, 7, 8, 8a, 8b and 13 |
| UFV-Foc590 | 2018 | Daniel Heck/ Eduardo Mizubuti | Teixeiras | Southeast | MG | -42,8337730 | -20,6384300 | 690 | Banana | Maca | AAB | - | NT | + | 1 |
| UFV-Foc592 | 2018 | Daniel Heck/ Eduardo Mizubuti | Vicosa | Southeast | MG | -42,8786000 | -20,7549000 | 690 | Banana | Maca | AAB | - | NT | + | - |
| UFV-Foc705 | - | - | Barbalha | Northeast | CE | -39,3025000 | -7,3055100 | 415 | Banana | Prata | AAB | B | 0124/22 | + | 1 and 13 |
| UFV-Foc706 | - | - | Barbalha | Northeast | CE | -39,3025000 | -7,3055100 | 415 | Banana | Prata | AAB | B | 0124/5/8/22 | + | 1 and 13 |
| UFV-Foc707 | - | - | Barbalha | Northeast | CE | -39,3025000 | -7,3055100 | 415 | Banana | Prata | AAB | B | 0124 | + | 1 and 13 |
| UFV-Foc708 | - | - | Barbalha | Northeast | CE | -39,3025000 | -7,3055100 | 415 | Banana | Prata | AAB | B | 0125/8/20 | + | 1 and 13 |
| UFV-Foc709 | - | - | Barbalha | Northeast | CE | -39,3025000 | -7,3055100 | 415 | Banana | Prata | AAB | B | 0124/5/22 | + | 1 and 13 |
| UFV-Foc711 | - | - | Barbalha | Northeast | CE | -39,3025000 | -7,3055100 | 415 | Banana | Prata | AAB | B | *crn* mutant | *+* | *-* |
| UFV-Foc712 | - | - | Barbalha | Northeast | CE | -39,3025000 | -7,3055100 | 415 | Banana | Prata | AAB | B |  | + | 1 and 13 |
| UFV-Foc713 | - | - | Barbalha | Northeast | CE | -39,3025000 | -7,3055100 | 415 | Banana | Prata | AAB | A | 0122 | + | - |
| UFV-Foc715 | - | - | Itinga do Maranhão | Northeast | MA | -47,5300000 | -4,4521500 | 144 | Banana | Prata | AAB | B | 0124/5/22 | + | 1 and 13 |
| UFV-Foc716 | - | - | Itinga do Maranhão | Northeast | MA | -47,5300000 | -4,4521500 | 144 | Banana | Prata | AAB | B | 0124/5/22 | + | 1 and 13 |
| UFV-Foc775 | 2017 | Jânia Bentes | Manaus | North | AM | -60,0261000 | -3,1071900 | 39 | Banana | - | - | - | Unknown | + | - |
| UFV-Foc776 | 2017 | Jânia Bentes | Manaus | North | AM | -60,0261000 | -3,1071900 | 39 | Banana | - | - | - | Unknown | + | 1, 8a, 8b and 13 |
| UFV-Foc777 | 2018 | Daniel Heck/ Eduardo Mizubuti | Teixeiras | Southeast | MG | -42,8262700 | -20,6314960 | 695 | Banana | Prata | AAB | - | NT | + | 1 and 8 |
| UFV-Foc785 | 2009 | Daniel Schurt | Boa Vista | North | RR | -60,6714000 | -2,8195400 | 76 | Banana | - | - | - | Unknown | + | - |
| UFV-Foc793 | - | Gilson Silva | São Luís | Northeast | MA | -44,3068000 | -2,5300000 | 4 | Banana | - | - | - | *crn* mutant | *+* | *-* |
| UFV-Foc795 | 2017 | Jânia Bentes | Manaus | North | AM | -60,0261000 | -3,1071900 | 39 | Banana | - | - | - | NT | + | - |
| UFV-Foc796 | 2017 | Jânia Bentes | Manaus | North | AM | -60,0261000 | -3,1071900 | 39 | Banana | - | - | - | Unknown | + | 1 and 13 |
| UFV-Foc797 | 2017 | Jânia Bentes | Manaus | North | AM | -60,0261000 | -3,1071900 | 39 | Banana | - | - | - | Unknown | + | - |
| UFV-Foc798 | 2017 | Jânia Bentes | Manaus | North | AM | -60,0261000 | -3,1071900 | 39 | Banana | - | - | - | Unknown | + | - |
| UFV-Foc799 | 2017 | Jânia Bentes | Manaus | North | AM | -60,0261000 | -3,1071900 | 39 | Banana | - | - | - | Unknown | + | 1 |

VCG: Vegetative compatibility group identified in this study

NT: Not tested

g: Genotypede isolates

**Supplementary table 2**  Primer sequences and PCR conditionsused for multiplex PCR analysis in this study.

| **Primer name** | **Sequence 5’-3’** | **Target gene** | **Size (bp)** | **Reference** | **PCR multiplex** |
| --- | --- | --- | --- | --- | --- |
| Six1-2 | confidential data | *confidential data* | *122* | Unpublished | M1 |
| Six1-2 | confidential data |  |  |  |  |
| FocTR4F | CACGTTTAAGGTGCCATGAGAG | *IGS* | *463* | Dita et al. (2010) | M1 |
| FocTR4R | GCCAGGACTGCCTCGTGA |  |  |  |  |
| Foc-1 | CAGGGGATGTATGAGGAGGCT | *OP02* | *242* | Lin et al. (2009) | M1 |
| Foc-2 | GTGACAGCGTCGTCTAGTTCC |  |  |  |  |
| SIX1-F | ATGGTACTCCTTGGCGCCCTC | *SIX1* | *260* | Meldrum et al. (2012) | M2 |
| SIX1-R | TGACAATGCGACCACGCCTCG |  |  |  |  |
| SIX2-F | ACGACCTGGGCCATCTCGGT | *SIX2* | *660* | Meldrum et al. (2012) | M2 |
| SIX2-R | ACACCTTGACTGCGACGCAACG |  |  |  |  |
| SIX3-F | ACCGACCATCTTGCCTAAACATTTACC | *SIX3* | *555* | Meldrum et al. (2012) | M3 |
| SIX3-R | TTAACCACTCTGCCAAGGGGAACT |  |  |  |  |
| SIX4-F | TGCTTCGGTGGCTGTTACATCTGC | *SIX4* | *808* | Meldrum et al. (2012) | M2 |
| SIX4-R | CCTAACCTAAGCTCACCCTCAGGAA |  |  |  |  |
| SIX5-F | TGCGCTTCGAGTACATCTCTGTTC | *SIX5* | *326* | Meldrum et al. (2012) | M2 |
| SIX5-R | CTGGTGAGATTTAGAGCAGTCAAAGCA |  |  |  |  |
| SIX6-F | GGCTGCGTAGCTGGTCCCCT | *SIX6* | *611* | Meldrum et al. (2012) | M5 |
| SIX6-R | CATGTCATGAATGTACGCATGTCCCT |  |  |  |  |
| SIX7-F | ACCTTTACCTCCTTTTCCATTTCGCCC | *SIX7* | *610* | Meldrum et al. (2012) | M3 |
| SIX7-R | CGAAAGTCAGCAAGGCCCCTGG |  |  |  |  |
| SIX8-F | TCGCCTGCATAACAGGTGCCG | *SIX8* | *250* | Meldrum et al. (2012) | M3 |
| SIX8-R | TTGTGTAGAAACTGGACAGTCGATGC |  |  |  |  |
| Foc-SIX8-F | CGAAGTGCGCCATATAAGACT | *Foc-SIX8a&b* | *770* | Fraser-Smith et al. (2014) | M4 |
| Foc-SIX8-R | CACCTGCTTGCTCCTTATCC |  |  |  |  |
| Foc-SIX8b-F | CGTCCTTACTTATATACCCTCTCAA | *Foc-SIX8b* | *595* | Fraser-Smith et al. (2014) | M4 |
| Foc-SIX8b-R | GGCCTAATCCACACAACA |  |  |  |  |
| SIX9-F2 | CTTCTAGCAGTTGTAGCCAC | *SIX9* | ~~319 | Schmidt et al. (2013) and Laurence et al. (2015) | M3 |
| SIX9-R2 | GTACGCCAGTTGACGCAAG |  |  |  |  |
| SIX10-F | AAAAAGCAGGCTCCATGAAGCTCTTGTGGTTG | *SIX10* | ~~550 | Schmidt et al. (2013) and Laurence et al. (2015) | M3 |
| SIX10-R | AGAAAGCTGGGTCCTACTTAGACCTGGTAATTGTT |  |  |  |  |
| S11f | GATGTTCTCCAAAGCCATCC | *SIX11* | *306* | Kashiwa et al. (2017) | Singleplex 1 |
| S11r | AGAATGCCACTCGGTGTGA |  |  |  |  |
| S12f | CAAGCGTCCAGTTGTCTCAG | *SIX12* | *312* | Kashiwa et al. (2017) | M5 |
| S12r | TCAGGAGTGGCATAGCTTGG |  |  |  |  |
| S13f | GATCAGGCCTTCAACGAAGA | *SIX13* | *738* | Kashiwa et al. (2017) | M5 |
| S13r | TAACTCGGCATCGATGGAAT |  |  |  |  |
| SIX14-F | TTGCCACCTATGCATACCG | *SIX14* | ~320 | Schmidt et al. (2013) and Laurence et al. (2015) | Singleplex 2 |
| SIX14-R | TCCACATTCCTAAGCGAACC |  |  |  |  |

**Supplementary table 3** Vegetative compatibility tester strains used in this study.

| **Code** | **Species** | **Formae speciales** | **Mutation** | **Cultivar** | **Location** | **Country** | **VCG** | **Provided by** | **Year** | **Race**^1^ |
| --- | --- | --- | --- | --- | --- | --- | --- | --- | --- | --- |
| O-1219-2 | *Fusarium oxysporum* | *cubense* | *nit*M | Mons | Queensland | Australia | 0120 | Randy Ploetz, UF | 2017 | 4? |
| O-1219-10 | *Fusarium oxysporum* | *cubense* | *nit*1 | Mons | Queensland | Australia | 0120 | Randy Ploetz, UF | 2017 | 4? |
| F9130-2 | *Fusarium oxysporum* | *cubense* | *nit*1 | Cavendish | - | Taiwan | 0121 | Randy Ploetz, UF | 2017 | 4 |
| F9130-7 | *Fusarium oxysporum* | *cubense* | *nit*M | Cavendish | - | Taiwan | 0121 | Randy Ploetz, UF | 2017 | 4 |
| PH2-3 | *Fusarium oxysporum* | *cubense* | *nit*1 | Cavendish | - | Philippines | 0122 | Randy Ploetz, UF | 2017 | 4? |
| PHL1-8 | *Fusarium oxysporum* | *cubense* | *nit*M | Latundan | - | Philippines | 0123 | Randy Ploetz, UF | 2017 | ? |
| PHL2-8 | *Fusarium oxysporum* | *cubense* | *nit*1 | Latundan | - | Philippines | 0123 | Randy Ploetz, UF | 2017 | ? |
| BLUG15 | *Fusarium oxysporum* | *cubense* | *nit*M | Bluggoe | - | Honduras | 0124 | Randy Ploetz, UF | 2017 | 2 |
| S?-1 | *Fusarium oxysporum* | *cubense* | *nit*1 | Tetraploid 1242 | Bodles | Jamaica | 0124 | Randy Ploetz, UF | 2017 | 2 |
| 8610-2 | *Fusarium oxysporum* | *cubense* | *nit*1 | Lady finger | Murwillumbah | Australia | 0125 | Randy Ploetz, UF | 2017 | 1 |
| 8611-9 | *Fusarium oxysporum* | *cubense* | *nit*M | Lady finger | Queensland | Australia | 0125 | Randy Ploetz, UF | 2017 | 1 |
| STM2-1 | *Fusarium oxysporum* | *cubense* | *nit*M | Maqueno | - | Honduras | 0126 | Randy Ploetz, UF | 2017 | 1 |
| 22994-5 | *Fusarium oxysporum* | *cubense* | *nit*1 | Bluggoe | South Johnstone | Australia | 0128 | Randy Ploetz, UF | 2017 | 2 |
| 22994-6 | *Fusarium oxysporum* | *cubense* | *nit*M | Bluggoe | South Johnstone | Australia | 0128 | Randy Ploetz, UF | 2017 | 2 |
| 8622-9 | *Fusarium oxysporum* | *cubense* | *nit*1 | Cavendish | Queensland | Australia | 0129 | Randy Ploetz, UF | 2017 | 4? |
| A1-1-7 | *Fusarium oxysporum* | *cubense* | *nit*1 | Apple | Florida | United State | 01210 | Randy Ploetz, UF | 2017 | 1? |
| A1-1-9 | *Fusarium oxysporum* | *cubense* | *nit*M | Apple | Florida | United State | 01210 | Randy Ploetz, UF | 2017 | 1? |
| SH3142-3 | *Fusarium oxysporum* | *cubense* | *nit*M | SH 3142 | Queensland | Australia | 01211 | Randy Ploetz, UF | 2017 | ? |
| SH3142-4 | *Fusarium oxysporum* | *cubense* | *nit*1 | SH 3142 | Queensland | Australia | 01211 | Randy Ploetz, UF | 2017 | ? |
| STNP2-3 | *Fusarium oxysporum* | *cubense* | *nit*M | Ney Poovan | Tenguero Station | Tanzania | 01212 | Randy Ploetz, UF | 2017 | ? |
| STNP4-8 | *Fusarium oxysporum* | *cubense* | *nit*1 | Ney Poovan | Bukaba Station | Tanzania | 01212 | Randy Ploetz, UF | 2017 | ? |
| MA2-2 | *Fusarium oxysporum* | *cubense* | *nit*M | Harare | Karonga | Malawi | 01214 | Randy Ploetz, UF | 2017 | ? |
| MW40-1 | *Fusarium oxysporum* | *cubense* | *nit*1 | Harare | Karonga | Malawi | 01214 | Randy Ploetz, UF | 2017 | ? |
| Cr2-3-1 | *Fusarium oxysporum* | *cubense* | *nit*M | Gros Michel | Isolona | Costa Rica | 01215 | Randy Ploetz, UF | 2017 | ? |
| Cr2-2-1 | *Fusarium oxysporum* | *cubense* | *nit*1 | Gros Michel | Hamburgo | Costa Rica | 01215 | Randy Ploetz, UF | 2017 | ? |
| Indo25-9 | *Fusarium oxysporum* | *cubense* | *nit*1 | Pisang Ambon | Sumatra | Indonesia | 01219 | Randy Ploetz, UF | 2017 | ? |
| 24223 | *Fusarium oxysporum* | *cubense* | *nit*M | Williams | Carnarvon | Australia | 01220 | Randy Ploetz, UF | 2017 | ? |
| RPML9-1 | *Fusarium oxysporum* | *cubense* | *nit*1 | Pisang awak legor | Pantai Acheh | Malaysia | 01222 | Randy Ploetz, UF | 2017 | ? |
| RPML4-6 | *Fusarium oxysporum* | *cubense* | *nit*M | Pisang awak legor | Pantai Acheh | Malaysia | 01222 | Randy Ploetz, UF | 2017 | ? |

^1^ Information provided by the collector.

**Supplementary table 4** Primer sequences, SSR motifs, number of alleles and allele sizes, used in this study.

| **SSR**  **Primer^2^** | **Locus** | **Motifs** | **Primers sequences 5′–3′** | **Fluorescent dye** | **Allele size range (bp)** | **No. of alleles^1^** |
| --- | --- | --- | --- | --- | --- | --- |
| MB2 | FO2 | (GT)11(GA)6 | F: TGCTGTGTATGGATGGATGG | NED | 240-272 | 10 |
|  |  |  | R: CATGGTCGATAGCTTGTCTCAG |  |  |  |
| MB5 | FO5 | (TG)9 | F: ACTTGGAGGAAATGGGCTTC | VIC | - | - |
|  |  |  | R: GGATGGCGTTTAATAAATCTGG |  |  |  |
| MB9 | FO9 | (CA)9 | F: TGGCTGGGATACTGTGTAATTG | NED | - | - |
|  |  |  | R: TTAGCTTCAGAGCCCTTTGG |  |  |  |
| MB10 | FO10 | (AAC)6 | F: TATCGAGTCCGGCTTCCAGAAC | 6FAM | - | - |
|  |  |  | R: TTGCAATTACCTCCGATACCAC |  |  |  |
| MB11 | FO11 | (GGC)7 | F: GTGGACGAACACCTGCATC | VIC | 173-182 | 4 |
|  |  |  | R: AGATCCTCCACCTCCACCTC |  |  |  |
| MB13 | FO13 | (CTTGGAAGTGGTAGCGG)14 | F: GGAGGATGAGCTCGATGAAG | PET | - | - |
|  |  |  | R: CTAAGCCTGCTACACCCTCG |  |  |  |
| MB14 | FO14 | (CCA)5 | F: CGTCTCTGAACCACCTTCATC | 6FAM | 181-184 | 2 |
|  |  |  | R: TTCCTCCGTCCATCCTGAC |  |  |  |
| MB17 | FO17 | (CA)21 | F: ACTGATTCACCGATCCTTGG | 6FAM | 302-334 | 10 |
|  |  |  | R: GCTGGCCTGACTTGTTATCG |  |  |  |
| MB18 | FO18 | (CAACA)6 | F: GGTAGGAAATGACGAAGCTGAC | PET | 254-294 | 6 |
|  |  |  | R: TGAGCACTCTAGCACTCCAAAC |  |  |  |

^1^Number of observed alleles.

^2^Simple sequence repeats primers (SSRs) described by Bogale et al. 2005.
